# Array Detection Enables Large Localization Range for Simple and Robust MINFLUX

**DOI:** 10.1101/2024.07.08.602588

**Authors:** Eli Slenders, Sanket Patil, Marcus Oliver Held, Alessandro Zunino, Giuseppe Vicidomini

## Abstract

The MINFLUX concept significantly enhances the spatial resolution of single-molecule localization microscopy (SMLM) by overcoming the limit imposed by the fluorophore’s photon counts. Typical MINFLUX microscopes localize the target molecule by scanning a zero-intensity focus around the molecule in a circular trajectory, with smaller trajectory diameters yielding lower localization uncertainties for a given number of photons. Since this approach requires the molecule to be within the scanned trajectory, MINFLUX typically relies on a photon-demanding iterative scheme with decreasing trajectory diameters. Although the iterative procedure does not substantially reduce the photon efficiency of MINFLUX, this approach is prone to misplacements of the trajectory and increases the system’s complexity. In this work, we introduce ISM-FLUX, a novel implementation of MINFLUX using image-scanning microscopy (ISM) with a single-photon avalanche diode (SPAD) array detector. ISM-FLUX provides precise MINFLUX localization within the trajectory while maintaining conventional photon-limited uncertainty outside it. The robustness of ISM-FLUX localization results in a larger localization range and greatly simplifies the architecture, which may facilitate broader adoption of MIN-FLUX.

## Introduction

Single-molecule localization microscopy (SMLM) achieves molecular resolution by sequentially localizing the fluorescent molecules that tag the structure of interest with nanometer precision (1). Original SMLM techniques, such as photoactivated localization microscopy (PALM) (2), stochastic optical reconstruction microscopy (STORM) (3), and points accumulation for imaging in nanoscale topography (PAINT) (4, 5), are based on simple widefield architectures. In these techniques, a relatively large area of the sample (hundreds of µm2) is uniformly illuminated by a laser source, while with a high quantum efficiency, the low-noise scientific camera registers a series of images. The sparsity of activated molecules within each camera image allows for their localization and subsequent reconstruction of a super-resolved image, whose resolution primarily depends on the localization uncertainty. Assuming negligible background, the localization uncertainty scales with 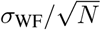, where *σ*_WF_ is the standard deviation of the widefield point-spread function, and *N* is the number of photons detected from the molecule during the camera frame time.

More recently, a new type of SMLM has been proposed that overcomes the traditional widefield-based approaches in terms of photon efficiency, *i*.*e*., the localization uncertainty for a given number of photons *N*. MIN-FLUX (6, 7) and derived techniques (8, 9) are based on a laser-scanning microscope whose focused excitation pattern features a zero-intensity point, typically an orbital angular momentum (OAM) beam. By scanning this focused beam around the molecule according to a precise targeted coordinate pattern (TCP) and analyzing the series of intensity values registered by a single-element detector during the scanning, it is possible to localize the molecule with an uncertainty that depends not only on the number of photons *N* but also on the size of the TCP. For a circular TCP of diameter *L*, the uncertainty scales as 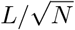. Notably, unlike *σ*_WF_, the parameter *L* is not bound by diffraction. As a result, these methods yield a localization uncertainty down to single-digit nanometers — and even below — using a limited number of photons, as typically provided by organic fluorophores. A potential advantage of MINFLUX is its compatibility with fluorescence lifetime analysis (8) due to the use of a single-photon detector, such as a single-photon avalanche diode (SPAD) detector.

However, a constraint of MINFLUX is that the molecule to be localized must reside within the TCP. If the molecule is outside the TCP, its position cannot always be uniquely determined from the data, and the localization uncertainty rapidly grows for a given number of photons (10). This limitation translates into a lack of robustness for the pure MINFLUX concept, which needs to be compensated by implementing iterative localization approaches. These approaches typically start with relatively large *L* values and a Gaussian beam illumination pattern before fully exploiting the benefits of MINFLUX by updating the TCP center after every iteration with a new estimate of the emitter position (7). The iterative implementation adds technical complexity to the otherwise straightforward laser-scanning microscope architecture. For example, rapidly moving the illumination minimum in an iterative TCP requires *e*.*g*., electro-optical devices, which have a limited range, and the iterative approach requires a precise estimate of the emitter position after every iteration.

Even though MINFLUX has shown superior results in single-molecule localization and tracking (11–13), the throughput and robustness might be further improved. We have recently theoretically shown that substituting the single-element detector with an array detector, such as an asynchronous read-out SPAD array detector, can maximize the localization range of MINFLUX (10). Asynchronous readout SPAD array detectors are transforming confocal laser-scanning microscopy (14–16). In contrast to single-element detectors, they provide an image of the laser-scanning microscope’s probed region. Initially, these properties allowed for a straightforward implementation of image-scanning microscopy (ISM), a technique that reduces the width of the PSF s of a laser-scanning microscope by a factor of 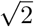 while maintaining a high signal-to-noise ratio (17–19). Another essential aspect of SPAD array detectors is that each element of the array can operate independently. As a result, asynchronous-readout SPAD array detectors achieve a temporal resolution comparable to conventional single-element detectors — without frame-rate limitations. The single-photon detection and timing ability allows correlation with fluorescence lifetime analysis (14, 20–23). More recently, asynchronous read-out SPAD array detectors have been used to implement an ISM-based SMLM technique (24). Assuming that the raster-scanning apparatus of the microscope is fast enough to provide true ISM images of the activated molecules in the sample, the dataset can be used as in widefield-based SMLM, with the advantage that the localization uncertainty is given by the ISM point-spread function, which is 2 times smaller than in widefield microscopy, and the combination with fluorescence lifetime is feasible. While this approach further demonstrates the potential of this class of detectors for SMLM, it does not highlight their benefits for MINFLUX.

Here, we demonstrate a MINFLUX system based on an asynchronous read-out SPAD array detector that addresses the limitations of the iterative approach. Our system effectively localizes molecules within the TCP with the expected MINFLUX localization uncertainty, while for molecules outside the TCP, our system achieves a localization uncertainty comparable to that of widefield-based SMLM. In particular, the localization range *R* is extended to several hundreds of nm, *i*.*e. R* » *L*. Here, we demonstrate that another significant advantage of array detectors is the flexibility to select alternative TCP configurations, such as circular patterns, with similar performance in terms of localization range and precision. This flexibility directly translates into a more simplified setup. Indeed, we experimentally validate this principle using an asynchronous read-out SPAD array detector installed in an ISM equipped with a standard galvanometer system for scanning, where only minor modifications have been introduced to shape the excitation focus as a doughnut. We call this method ISM-FLUX because it leverages the capability of the SPAD array to image the probed region of a laser scanning microscope rather than operating on the reconstructed ISM image. This aspect allows us to localize molecules with the photon efficiency of MINFLUX and not be limited to a 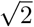 factor.

## Results

The ISM-FLUX setup is based on a conventional confocal microscope with only two modifications (Fig. 1) (a). Firstly, a vortex phase plate conjugated with the objective back focal plane is installed in the illumination path to generate a doughnut-shaped excitation focus. Secondly, instead of a single-element detector, a 5×5 SPAD array detector is installed in the image plane in descanned mode. A 3D active stabilization system based on tracking fiducial markers keeps the sample in place with about 1-2 nm precision (standard deviation) for all three axes.

**Fig. 1.**
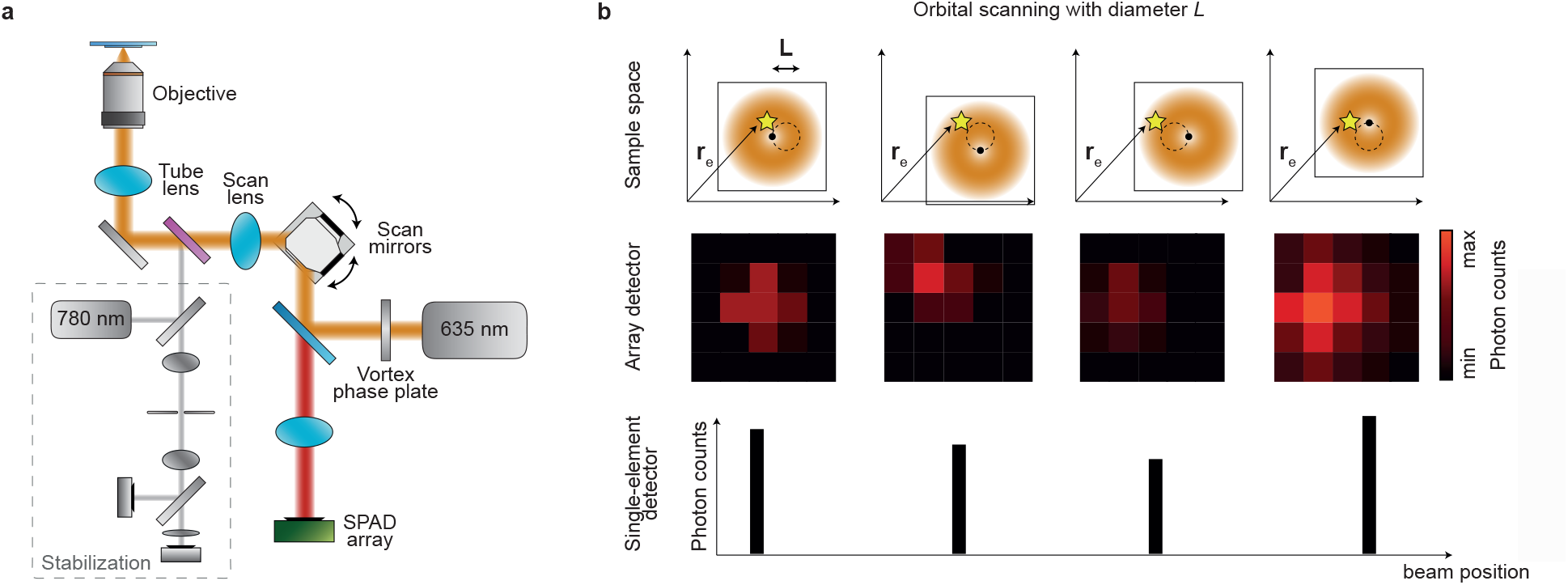
Concept of single-molecule localization with ISM-FLUX. (a) The ISM-FLUX setup is based on a confocal microscope equipped with a vortex phase plate to obtain a doughnut-shaped excitation beam and with a SPAD array detector. A 3D stabilization system based on tracking fiducial markers keeps the sample steady, omitting the need for drift correction in post-processing. A detailed scheme of the setup and the stabilization unit is shown in (Fig. S10). (b) The doughnut-shaped excitation beam moves in a circular pattern around a single emitter (yellow star). For each position on the circle, the array detector registers a micro-image of the emitted fluorescence. The final photon count data set is four-dimensional: *I*(*c, α, x*_*d*_, *y*_*d*_), with *c* the orbit number, *α* the angular coordinate within the orbit, and (*x*_*d*_, *y*_*d*_) the detector element coordinates of the array detector. For simplicity, we describe each detector element with a number *I*(*c, α, d*). In conventional MINFLUX-based techniques, a single-element detector counts the number of fluorescence photons, discarding their position in the image plane, *I*(*c, α*).

The TCP was generated by using the galvanometric scan mirrors to steer the doughnut beam to 32 positions on a circle with a diameter of about *L* = 90 nm at a rate of 1.92 ms/circle (∼521 Hz) (Fig. 1) (b). For each position on the circle, an image of the probed sample region is collected – the so-called microimage. When a single emitter is present, these images vary in two aspects: the total number of detected photons per image and the spatial distribution of photons across the detector, which traces a circular path due to the descanned detection mode. While MINFLUX localizes solely based on photon counts, ISM-FLUX additionally considers the spatial coordinate of the detection event.

The usable information for position estimation of an emitter is summarized in an event data set of *N*_*c*_×*N*_*d*_×*N*_*t*_ photon counts, with *N*_*c*_ the number of positions in the circle (*N*_*c*_ = 32), *N*_*d*_ the number of detector elements (*N*_*d*_ = 25), and *N*_*t*_ the number of consecutive orbits for which photons of the emitter were detected. Summing over the *N*_*t*_ orbits, we have a reduced event data set *n*_*ij*_, with *I* ∈ {1, 2, …, *N*_*c*_} and *j* ∈ {1, 2, …, *N*_*d*_}.

To find the emitter position, we extended the maximum likelihood approach from MINFLUX to the use of an array detector (10). The likelihood function for the emitter at position **r**_*E*_ can be written as

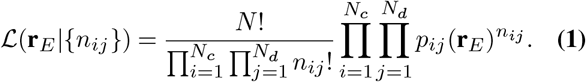

where, *p*_*ij*_(**r**_*E*_) are the molecule detection functions (MDFs), *i*.*e*., the emitter position-dependent probability to detect a photon during the *i*-th position in the circle and in the *j*-th element of the detector. All MDFs need to be known for the localization and can either be simulated using Eq. S18 or experimentally measured by raster-scanning an emitter.

The minimal uncertainty *σ*_CRB_ in localizing an emitter with an unbiased estimator, *i*.*e*., the Cramér–Rao bound, depends on various factors, such as *L, N*, and the position of the emitter relative to the center of the TCP. For an emitter at the center of the scanned orbit, 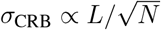 (10) (Fig. S3). Thus, the smaller the diameter of the orbital scan, *i*.*e*., the lower the *L* value, the more precisely the emitter can be localized for a given *N*. For a fixed *L* value, *σ*_CRB_ scales with 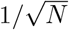. These properties are shared by both MINFLUX and ISM-FLUX. Similar properties hold for positions in close proximity to the TCP center. In this scenario, the emitter’s position is primarily inferred from the total number of photons detected at each position within the TCP, with the photon distribution on the array detector contributing minimal additional information.

However, a key difference between both techniques emerges when the emitter is located outside the circle described by the TCP. With a single-element detector, the localization range is restricted to a circle with a diameter below the TCP diameter. Outside of this range, *σ*_CRB_ diverges (Fig. 2) and the MLE is highly biased (Fig. S2). In contrast, the MLE with array detectors shows a non-diverging *σ*_CRB_ and an unbiased position estimation, at least for high photon counts, for a much larger range (Fig. 2) (a,b), (Fig. S2). Farther from the TCP center, the dominant source of information shifts to the image of the emitter, resulting in an increased *σ*_CRB_ that approaches the value of camera-based techniques. As a result, the localization range is not set by *L* but by the FOV of the detector, with a *σ*_CRB_ that varies depending on the emitter’s position. *E*.*g*., for *L* = 90 nm and 100 detected photons, *σ*_CRB_ increases from about 3.5 nm in the center to about 12 nm near the edge of the detector. The larger *L*, the more homogeneous *σ*_CRB_ becomes.

**Fig. 2.**
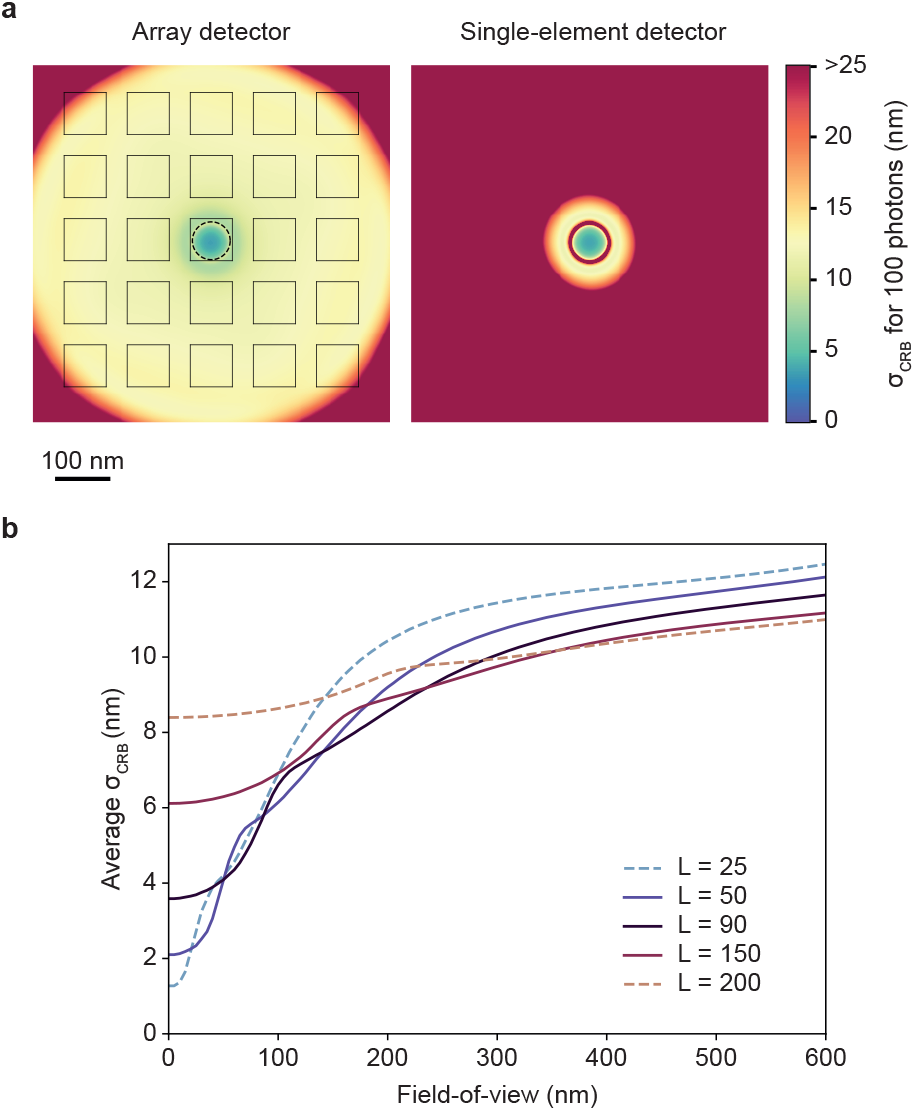
CRB as a function of the emitter position for 100 detected photons. (a) Comparison between an array detector and a single-element detector for *L* = 90 nm. (b) Average CRB within a circular FOV as a function of the FOV diameter. FOV=0 corresponds to the TCP center, and FOV=*L* to the TCP circle. Comparison of the CRB for different *L* values.

In our implementation, we kept the laser power, the TCP position, and the *L* value constant during the localization process. As a consequence, also the emission rate of the emitter depends on its position with respect to the TCP (Fig. S4). Due to the varying *N*, the effective localization uncertainty within the localization range is between 3.5-7 nm for *L* =90 nm and a photon count between 100 and 650. We chose a constant *L* value of 90 nm as a trade-off between minimum localization uncertainty in the center and maximum *σ*_CRB_ homogeneity. Note that keeping *L* large enough also keeps the difference between the expected photon counts for an emitter in the TCP center and an emitter near the doughnut maximum below one order of magnitude (Fig. S4). Since we do not adjust the laser power depending on the emitter’s position, the radiant flux received by a molecule in the TCP center should be sufficiently high to detect its fluorescence signal above the background. Simultaneously, for a molecule positioned near the peak of the doughnut-shaped illumination, the radiant flux should be low enough not to induce blinking, bleaching, or saturation of the fluorescence.

Note that we discretized the TCP orbit into 32 points. The calculation of *σ*_CRB_ shows that very similar results can be obtained with fewer probed points (Fig. S5). However, to ensure a smooth orbital scan in which potential inertia effects caused by the galvo’s motion and stopping is minimal, we opted to use 32 points.

### Enhanced localization range with ISM-FLUX

We validated the localization performance of ISM-FLUX by repeatedly localizing a gold nanoparticle (GNP) translated to different positions with a nanometer-precise piezo-electric stage. During the entire temporal series, the particle moved to different positions, the TCP remained constant, and the SPAD array detected photons in synchronization with the galvanometer mirror. We began by checking the setup’s performance in localizing a particle positioned inside or near the TCP circle (Fig. 3) (a). We moved the particle to 17 different positions, forming the logo of the Italian Institute of Technology (IIT). We calculated the localization uncertainty using 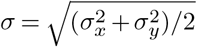, with *σ*_*x*_ and *σ*_*y*_ being the standard deviations of the estimated positions in *x* and *y*, respectively. For the three positions closest to the TCP center, *σ* is below 4 nm for an average of 364 detected photons (Fig. 3) (c). To evaluate position estimation with different photon counts, we binned the temporal series in post-processing for increasing numbers of TCP, resulting in a continuously decreasing localization uncertainty, which approached a limiting value of about 2 nm for 3000 photons. In addition to the uncertainty arising from photon statistics, other factors, such as residual vibrations and system drift, contribute to the overall uncertainty limit.

**Fig. 3.**
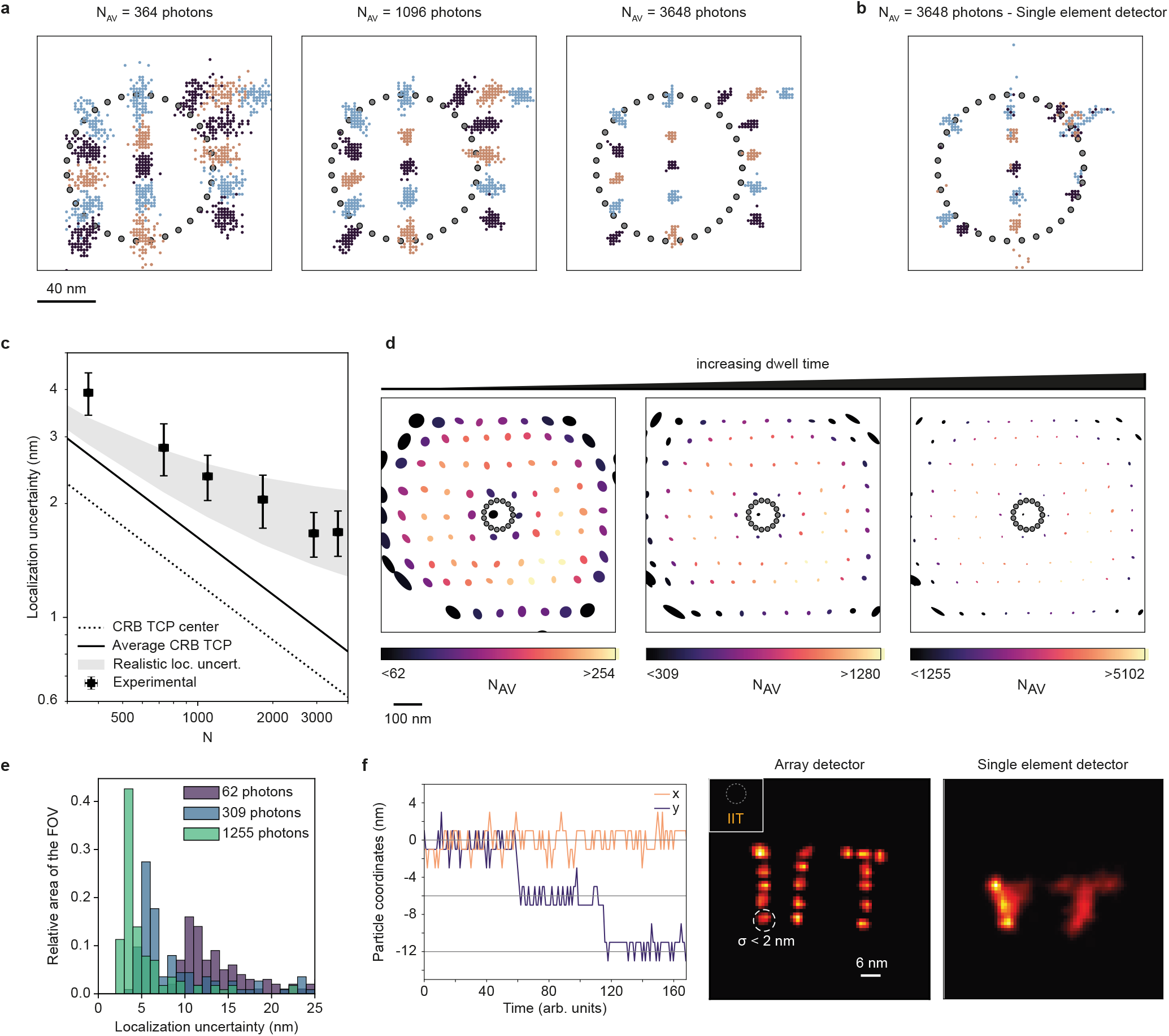
ISM-FLUX on a GNP. (a) Retrieved localizations for different particle positions were obtained by moving the piezo-stage to 17 different positions and different photon numbers. (b) Treating the data as if they were recorded with a single-element detector. (c) Average localization uncertainty of ISM-FLUX as a function of *N*, calculated as the average and standard deviation for the 3 positions in the pattern closest to the TCP center. The comparison with *σ*_CRB_ for the TCP center (dotted line) and the average within the TCP (solid line) are plotted. The ‘realistic localization uncertainty’ is calculated as 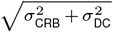, with *σ*_DC_ = 1 − 2 nm. (d) Calculated localization uncertainty over a large FOV, obtained by moving the GNP to 170 positions. The color shows the number of detected photon counts for a constant dwell time. The minimum color values (*i*.*e*., 62, 309, 1255) correspond to the photon counts for the TCP center. Close to the FOV edge, the actual number of photons may be lower, as indicated by the *<*-sign in the color bar. The size of the ellipses represents the localization uncertainty along two axes, see Methods. (e) Histograms of the data from (d), in which the localization uncertainty is defined as the average of half of the width and height of the ellipses. The relative area of the FOV is approximated as the number of positions in (d) divided by the total number of positions to which we moved the nanoparticle. The number of photons in the legend refers to the number of photons at the doughnut center; the actual number scales with increasing illumination, as indicated in (d). (f) Our ISM-FLUX system has a minimum localization uncertainty of about 2 nm, limited by the stabilization performance and computational power needed for the MLE. For high enough photon counts, we reach this limit of *σ <* 2 nm, and, as a result, we can resolve a pattern with 6 nm steps. The time trace shows a portion of the calculated (*x, y*) coordinates as a function of time. The images show the reconstruction for all particle positions and a comparison obtained by treating the data as if they were recorded with a single-element detector. The GNP is outside the TCP for all positions. The experimental MDFs were used for analysis. See (Fig. S6) for localization results of (a) and (f) with the simulated MDFs.

For comparison, we mimicked the scenario of a single-element detector by summing all photon counts across the 25 elements of the array detector. Estimating the position based on only 32 photon counts leads to a similar localization uncertainty for positions close to the TCP center (Fig. 3) (b). However, close to the border of the TCP and outside the TCP, we get an imprecise and biased estimate of the particle, underlining the limited localization range with a single-element detector. Note that the virtual single-element detector and the array detector are the same size (1.4 Airy Units) and, therefore, have the same FOV. The increased localization range of ISM-FLUX is entirely due to the subdiffractional spatial sampling provided by the array detector.

To verify whether the localization range closely matches the FOV of the array detector, we moved the particle in a grid pattern across the entire array detector FOV (about 600×600 nm^2^) with a step size of 100 nm. At each grid position, we localized the particle after a predetermined number of orbits, leading to a different total number of detected photons (up to a factor of 4) per imposed position. Despite a varying number of photon counts, we observed a uniform *σ* across the FOV of the detector (Fig. 3) (d). This uniformity is maintained for longer integration times, resulting in a lower *σ* for all positions, according to 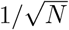 (Fig. 3) (d,e).

Building on these results, it remains to be shown that we can use the single-digit localization uncertainty outside the TCP to resolve single-digit spatial increments. Therefore, we repeated the experiment of GNP localization with a step size of 6 nm. For the GNP placed outside the TCP, we observed a *σ* value below 2 nm, and consequently, we could resolve the 6 nm steps (Fig. 3) (f). The ability to resolve these steps stems mainly from the subdiffractional sampling of the detected photons. Instead, a localization scheme based on a single-element detector would require additional prelocalization.

### Resolving DNA-origami nanorulers with ISM-FLUX

For high enough photon counts, we achieved single-digit localization uncertainty for bright scatterers. Here, we demonstrate the ability of ISM-FLUX to localize single fluorophores.

First, we used ISM-FLUX to localize fixed ATTO647N fluorophores. The sample was raster-scanned to find single fluorophores (Fig. 4) (a), followed by ISM-FLUX measurements at different positions close to the fluorophore positions (Fig. 4) (b,c). For a fluorophore in the TCP center, we found localization uncertainty values close to *σ*_CRB_ for all *N* values. Analyzing the same data for a single-element detector gives almost identical results. For a molecule outside the TCP, ISM-FLUX is less photo-efficient, as illustrated by the higher localization uncertainty. However, the spatial information from the array detector still results in an unbiased position estimation. Instead, with a single-element detector, we cannot retrieve the fluorophore position.

**Fig. 4.**
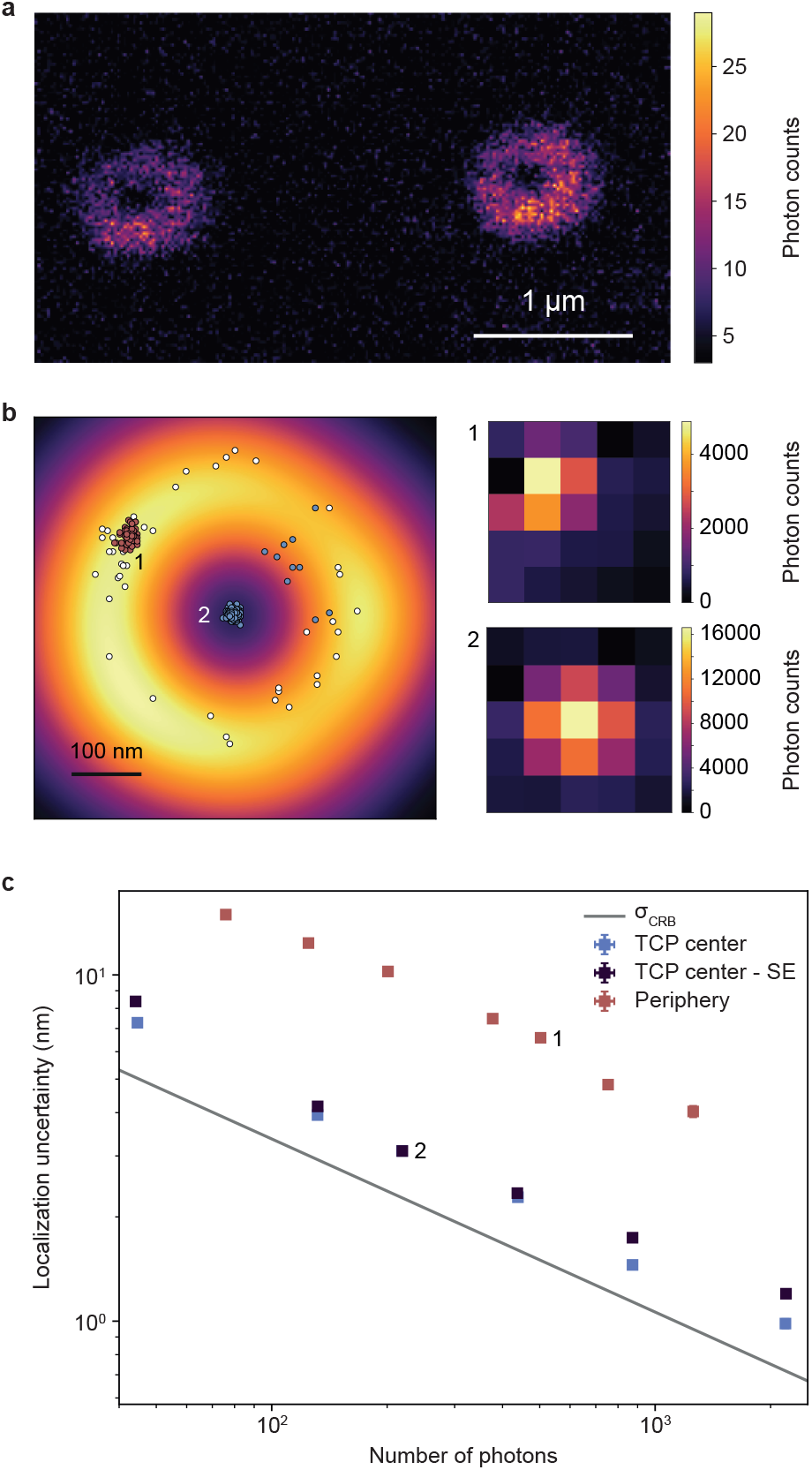
ISM-FLUX on fixed single ATTO647N molecules. (a) Raster-scanned image of individual fluorescent molecules, obtained by summing the signals of all detector elements for each scan position. (b) ISM-FLUX on two emitters, one in the periphery (1) and another in the TCP center (2), are sequentially localized until bleached. The white scatter plot represents the same data for the peripheral molecule, analyzed as if using a single-element detector. The fingerprints show the summed photon counts over all TCP positions and orbits. (c) Corresponding ISM-FLUX localization uncertainty for different photon counts, obtained by chopping the time trace in segments of different sizes. The two numbered data points correspond to the scatter plots in (a). SE = analysis as if a single-element detector was used. Error bars represent standard errors but are mostly too small to be visible. *σ*_CRB_ shows the Cramér–Rao bound for the TCP center.

Having demonstrated the increased localization range of ISM-FLUX on single molecules, we now demonstrate the resolving capability of ISM-FLUX with two types of DNA origami nanorulers with labeling distances of 20 nm and 40 nm. As a switching mechanism, we use DNA-PAINT, but ISM-FLUX is also compatible with other mechanisms.

We first took advantage of the confocal architecture by performing conventional raster-scanned imaging over a large field of view (FOV) (> 20 × 20 µm) to scout the sample and select origami positions. The large localization range of ISM-FLUX (about 600 × 600 nm^2^) eliminates the requirement for a more precise pre-localization step. For ISM-FLUX, we used an orbital scan frequency of 1.92 ms/orbit, which ensured that we could sum over many orbits for the duration of an average event time of 100 ms.

Given the ISM-FLUX measurement, ON-events were identified by thresholding the intensity trace (Fig. 5), and the following filters were applied to the on-events: (i) a lower threshold on the total number of photons of the event, (ii) an upper threshold on the intensity fluctuations during the on-event, and (iii) a lower threshold on the duration, see Methods.

**Fig. 5.**
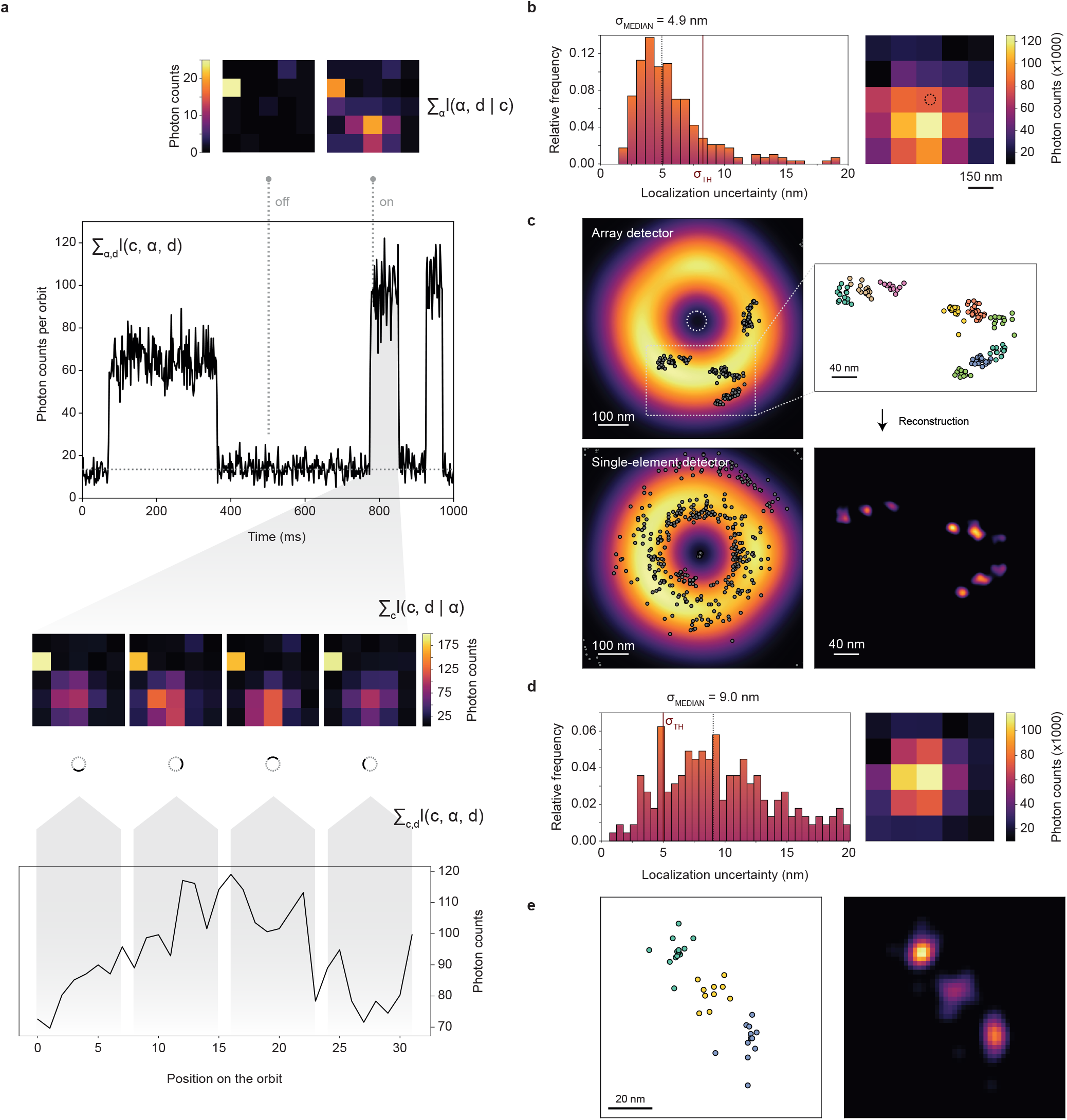
ISM-FLUX on DNA-origami nanorulers. (a-c) ISM-FLUX on DNA-origami with 40 nm spacing between nearest neighbors. (a) Different representations of the same data set show the on-off blinking and the signal on the detector for a representative on-event. (b) Histogram of the experimental localization uncertainty of all blinking events and the sum of all micro-images during the on-events. It shows a segment of a 9.6-minute ISM-FLUX measurement. (c) Scatter plot of the localizations (top-left). Only localizations that pass through all filters are shown. The background image shows the sums of all MDFs. The colored scatter plot (right) shows a zoomed-in region, with the three docking sites of each origami shown in a different color. Classification based on k-means clustering with 9 clusters. Reconstructed images made with a Gaussian kernel with a fixed standard deviation of 6 nm. The single-element detector plot (bottom-left) shows the localizations obtained by summing the signal over all detector elements, thus ignoring the spatial information on the detection side. (d-e) Data are similar to (b-c) for a DNA-origami sample with 20 nm spacing between nearest neighbors.

Each remaining event was split into three equal-duration segments and localized using the maximum likelihood approach. We calculated the experimental localization uncertainty *σ* for each event, resulting in a histogram (Fig. 5) (b), with a median value of 4.9 nm. Note that since each event is split into 3 subevents, the value of 4.9 nm corresponds to the localization uncertainty obtained for minimum 2000*/*3 ≈ 667 photons, of which only about 80% (534 photons) are signal and the remaining counts are either background or dark counts. We discarded events for which *σ* exceeded a threshold of 8.25 nm. For the remaining events, each set of three localizations was averaged, leading to a final result of 215 events. Summing the micro-images of all events, (Fig. 5) (b) the so-called “fingerprint” reveals the presence of at least one origami outside the TCP. However, plotting the individual localizations and the reconstructed image, (Fig. 5) (c), shows four origami, all located outside of the TCP. Thanks to the large localization range of ISM-FLUX, all origami can be viewed in a single measurement. Conversely, when analyzing the same dataset for a single-element detector, we see a highly biased and unprecise localization map, as predicted by the estimation theory.

We repeated the imaging and analysis procedure for the origami structure with 20 nm spacing, (Fig. 5) (d,e). As shown by the fingerprint, the origami is closer to the TCP but still outside the TCP. Because of small differences between the experimental and simulated MDFs close to the doughnut minimum caused by small aberrations in the imaging system, we used the experimental MDFs for the analysis of this origami. As a result, the localization uncertainty histogram shows a wider distribution, (Fig. 5) (e), compared to (Fig. 5) (c), for which we used the simulated MDFs. We set the threshold for accepting events to *σ <* 5 nm, resulting in 36 localizations for the three spots combined. The scatter plot and the reconstruction show that the 20 nm spacing between the docking sites can be resolved (Fig. 5) (e).

## Discussion

We demonstrated that ISM-FLUX can localize a single fluorescent molecule with an uncertainty between 2 to 10 nm for approximately 1000 photons and an orbital scanning diameter *L* of 90 nm. Thanks to the unique spatial information leveraged by the array detector, we achieved this localization uncertainty within a 600×600 nm^2^ region. Specifically, the localization uncertainty depends on the position of the fluorophore relative to the TCP, which is the lowest at the TCP center. Different from MINFLUX, the localization uncertainty in ISM-FLUX does not diverge for fluorophores outside of the TCP, resulting in greater robustness and an extended localization range. As a result, the typical iterative procedure of MINFLUX in which the position and diameter of the TCP are changed is not crucial. In addition, the array detector gives more flexibility in the choice of TCP. Unlike MINFLUX, probing the central TCP point is not required to have an unbiased MLE. The ability to have a pure circular TCP is advantageous because a galvanometric scanning system of a confocal microscope can perform orbital scanning at high speeds (25, 26). Due to various factors such as inertia, the actual scan diameter performed by the galvanometric scan system is lower than the diameter as imposed by the signal delivered to the galvanometer drivers by the microscope control software. For that reason, a calibration step to derive the exact orbit parameters is crucial for the ISM-FLUX data analysis. However, having an array detector in de-scanned mode greatly simplifies this procedure, as this configuration allows self-calibration (Sup. Note 3) and (Fig. S9).

A limitation of ISM-FLUX and MINFLUX is their lower localization throughput compared to camera-based SMLM due to the sequential single-molecule localization inherent in these approaches. Shorter fluorophore on-times and higher rates of on-events could partially speed up acquisition. In DNA-PAINT, a higher rate of on-events can be achieved by increasing the imager concentration, but this would significantly raise background noise. Alternatively, other labeling strategies can be explored, ranging from common stochastically photoswitchable dyes and fluorogenic dyes to newly developed photoactivatable dyes (27). In this context, ISM-FLUX, by using an array detector, opens up the possibility of localizing multiple fluorophores simultaneously (10), which can further increase throughput. A current technical drawback of ISM-FLUX is the lower photon-collection efficiency of SPAD array detectors compared to the single-element SPAD detectors typically used in MINFLUX, particularly when using far-red fluorophores. However, we predict that this limitation will soon be overcome with the introduction of red-enhanced SPAD array detectors (16, 28).

In this ISM-FLUX implementation, single-molecule localization is limited to two dimensions, but an extension to 3D is conceivable in several ways: encoding the axial information (i) on the detection side, (ii) on the excitation side, or (iii) in the fluorescence lifetime. In the first approach, the axial position can be encoded in the molecule image by placing a cylindrical lens in front of the array detector, as was shown in (29) for tracking and commonly used in camera-based localization. This approach is technically simple but does not leverage the benefits of the MINFLUX principle along the axial dimension. In the second approach, one can generate an intensity minimum in the axial direction on the excitation side and translate this along the third dimension, similar to 3D MINFLUX (7). Lastly, the axial position can be encoded in the fluorescence lifetime when combined with metal-induced energy transfer (MIET): the reduction of the molecule’s lifetime when it is in proximity to a metal film can be measured to determine its axial position. However, this method restricts localization to a thin layer near the metal film.

While ISM-FLUX does not require an iterative procedure like MINFLUX, it could still benefit from one. Specifically, a two-step ISM-FLUX implementation can be envisioned: the first step coarsely localizes the molecule within a 600×600 nm^2^ region, and in the second step, the TCP is centered on the molecule to fully leverage the MINFLUX concept. This implementation would also guarantee consistent localization uncertainty across the entire localization range of ISM-FLUX. An advantage of this implementation is its natural extension to real-time single-molecule tracking. In this scenario, another benefit of the large localization range of ISM-FLUX is that an incorrect real-time estimate of the emitter position, and consequently an incorrect updated TCP position, will not result in a failed localization. Instead, it will only lead to a suboptimal localization in terms of photon efficiency.

In summary, from an optical architecture perspective, converting a confocal laser scanning microscope into an ISM-FLUX system requires only the addition of a phase plate or a spatial light modulator and an array detector. From the microscope control perspective, circular scanning is necessary instead of conventional raster scanning. Combined with the greater penetration depth compared to total internal reflection-based techniques, the self-calibration procedure, and automated analysis, we foresee a future where ISM-FLUX enables the easy integration of SMLM with single-digit nanometer localization uncertainty into existing confocal microscopes. The impact of ISM-FLUX will further increase with its natural evolution: integrating photon-time tagging measurements to fully explore the emerging photon-resolved microscopy paradigm (20, 31).

## Methods

### A. Optical setup

The setup is based on a conventional ISM setup with some minor modifications. A continuous wave 635 nm wavelength laser beam (LDH-D-C-635M connected to a PDL 800-D driver, PicoQuant GmbH, Berlin, Germany) was cleaned up by coupling the beam into a single-mode polarizing maintaining fiber after passing through a Glan-Thompson Polarizer, a half-wave plate, and a 2-to-1 telescope to demagnify the beam. The output from the fiber was collimated and passed through a quarter-wave plate and halfwave plate (500-900 nm achr, B. Halle, Berlin, Germany) to generate circularly polarized light. The beam passed through a vortex phase plate (V-633-10-1, Vortex Lens for 633 nm, m=1, Vortex Photonics, Planegg, Germany) to generate a doughnut-shaped excitation beam, was reflected by a 643 nm dichroic beam splitter (F48-643, AHF, Germany) and was scanned with a set of two galvanometric scan mirrors (Saturn 1B 56S, ScannerMAX, Sanford, Florida, USA). The scanning system was coupled to a 60 mm Nikon scan lens and a 200 mm Nikon tube lens. All measurements were performed with a Nikon 100 × 1.4 oil objective (Nikon Plan Apo VC 100x/1.40 Oil OFN25 DIC N2). The laser power was set to 51 µW and 77 µW, measured in the sample plane, for the 20 nm and 40 nm origami, respectively. The fluorescence was collected in descanned mode, passing through the dichroic, a set of three filters (BSP01-785R-25 short-pass 785 nm, notch 642 nm, and LP01-633R-25 633 nm long-pass filter) and focused with a 300 mm lens (AC254-300-A-ML, ThorLabs, Germany), conjugated with the scan lens, onto a SPAD array detector (PRISM Light, Genoa Instruments, Genoa, Italy). The resulting FOV of the detector was about 1.5 Airy units, which meant that the detector also acted as a pinhole to remove the out-of-focus fluorescence background.

The sample was stabilized throughout the entire DNA-origami data acquisition using a feedback tracking system based on fiducial markers. Specifically, the stabilization system employs a near-infrared laser to illuminate gold particles—acting as fiducial markers—using a highly inclined and laminated optical sheet (HILO) configuration (32). Two images were captured using two independent cameras to localize the fiducial markers in three dimensions. A cylindrical lens was used to introduce astigmatism in one camera image to localize the particles axially. The localization coordinates were used to control a high-resolution three-dimensional piezo stage in real time, keeping the particles stationary with respect to the objective. A detailed scheme of the setup and the stabilization unit is shown in (Fig. S10).

### B. Electronics and software

The scan mirror drivers and SPAD array detector were connected to an FPGA module (NI PXIe-1071 with an NI PXIe-7856R board, National Instruments, USA) that was controlled with the BrightEyes-MCS software for imaging and custom-built LabVIEW software for circular scanning. Only the central 5×5 detector elements were read. A 3-axes piezo-electric stage (P-545.3R8H PInano XYZ, Physik Instrumente, Karlsruhe, Germany) was connected to an E-727 digital piezo controller (Physik Instrumente) and connected to a PC. The same PC (Dell Precision 5820 Tower X-Series) was used for data acquisition and controlling the sample stabilization).

### C. Sample Preparation

The gold nanoparticle sample was prepared using 100 nm diameter, OD 1 gold nanoparticles (Sigma Aldrich, Product Number: 742031). The gold nanoparticle suspension was sonicated for 15 minutes. Then, the solution was drop-cast on a coverslip previously coated with Poly-L-Lysine (Sigma Aldrich, Product Number: P4832) by incubating the coverslip with Poly-L-Lysine for 30 minutes at room temperature and washed with *ddH*_2_*O* afterward.

The sample for single fluorophore measurements contained passively adsorbed antibodies on an untreated glass surface in an 80% glycerol concentration. The stock solution of anti-mouse IgG-Atto647N (50185-1ML-F, Sigma Aldrich, Steinheim, Germany), was initially diluted in PBS, and for the final single-molecule concentration in a home-made 80% (w/w) glycerol solution (G5516-500ML, Sigma Aldrich, Steinheim, Germany). A coverslip of the dimension of a microscope slide (24 x 60 mm, No. 1.5H, Paul Marienfeld GmbH, Lauda-Königshofen, Germany) was cleaned with mQ water and mounted into the microscope. Right before the start of the measurement, 20ul of the antibody-glycerol solution was dropped onto the coverslip. Immediately afterward, antibodies that were adsorbed or moving on the coverslip were visible, as well as in a solution. Measurements were performed only on non-moving antibodies. To evaluate the single fluorophore signal, only the signal from the last intensity step before complete photobleaching was evaluated.

The DNA-Origami, with DNA-PAINT labeling, was commercially purchased from GATTAquant (GATTAquant GmbH, Munich, Germany). Both nanorulers consisted of three collinear spots with 40 nm spacings ((GATTA-PAINT HiRes NP 40) or 20 nm spacings (GATTA-PAINT HiRes NP 20R), and are labeled with Atto655. The average binding time (on-time) of the origami was 100 ms.

### D. Measuring the MDFs

ISM-FLUX data analysis requires a set of *N*_*c*_ ×*N*_*d*_ MDFs, with *N*_*c*_ the number of illumination positions and *N*_*d*_ the number of detector elements in the array. The MDF describes the position-dependent probability to excite and detect a fluorescence photon from a molecule at a given position within the sample for a given position in the TCP and a given detector element (33). We measured the MDFs in two steps. First, we measured *I*_0*j*_(**r**_*E*_) by scanning over a single GNP with the galvo scan mirrors and detecting the signal with the SPAD array detector, (Fig. S11) (a). The scan settings were FOV 2×2 µm2, pixel size 2 nm^2^, and a scan speed corresponding to 15-30 s per image. We averaged over 5-6 images while the stabilization software kept the particle in place. The laser power was adjusted to keep the total detected PCR (sum 5×5) around 1 MHz in the doughnut maximum. The 2 nm pixel size was chosen as a trade-off between better localization uncertainty (with a small pixel size) and smaller MDF data size (with a bigger pixel size). Due to small day-to-day changes in the experimental setup, we measured *I*_0*j*_(**r**_*E*_) before each series of experiments. *I*_0*j*_(**r**_*E*_) takes the form of 1000× px 1000 px×25 detector elements. We noted that for small FOVs, the images were stretched in the x-direction (the fast scan axis) due to the limited precision of the galvo scanner, (Fig. S11) (a). Therefore, we squeezed the images 20% in the x-direction in post-processing using the resize function from the scikit-image package in Python. The second step is using *I*_0*j*_(**r**_*E*_) to generate *I*_*ij*_(**r**_*E*_) by using the orbital scan calibration. Knowing the radius and starting angle or the orbit and the direction of rotation, we shifted the 25 images *I*_0*j*_(**r**_*E*_) in post-processing to each of the 32 scan positions, resulting in a full MDF data set, containing 1000 px ×1000 px ×800 images, (Fig. S11) (b). For measurements in which the two hot detector elements were excluded, the MDF data set contained 736 images. All images were smoothed with a combination of a 2D uniform filter from SciPy and a GaussianBlur from OpenCV-Python until visually smooth.

### E. Simulation of the MDFs and calculation of the CRB

To simulate *I*_0*j*_(**r**_*E*_), we used the s^2^ISM (34) and BrightEyes-ISM (35) Python packages. First, we acquired a conventional ISM dataset of a fixed HeLa cell stained for *α*-tubulin (details in (20)). The experimental shift vectors were obtained through phase correlation of the images in (a) with the image from the central detector element (12) (Fig. S12). We assumed no circular symmetry-breaking aberrations and a detector pitch smaller than one airy unit to ensure evenly-spaced shift vectors. From the shift vectors of the images generated by the inner 3×3 detector elements, we fitted the orientation, rotation, and magnification of the system. Using the fitted parameters, we simulated a set of 25 MDFs assuming 635 nm excitation wavelength, 660 nm emission wavelength, NA 1.4, refractive index 1.5, vortex phase plate as a mask (angular momentum 1), pixel pitch and pixel size of the 5×5 array detector projected into the sample space 150 nm and 100 nm, respectively. The simulation space pixel size was 2.5 nm, 1000×1000 pixels. To obtain *I*_*ij*_(**r**_*E*_), (Fig. S11) (b), we followed the same protocol as before, without the smoothing.

For the calculation of *σ*_CRB_, we used the code described in (10), using as input parameters the simulated MDFs, assuming a total of 100 detected photons at each position and SBR set to 10 to simulate optimal conditions. For the simulations of (Fig. S4), we scaled *σ*_CRB_ with 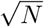, with *N* the expected number of photons at each position. The map of *N* (*x, y*) was found by summing all MDFs, rescaled to have *N* = 100 in the center.

### F. Measuring the orbit parameters and self-calibration procedure

The three parameters describing the spatial characteristics of the orbital scan are the diameter *L*, the starting phase *α*_0_, and the direction of rotation. All parameters may be derived from a reference measurement or from the SM measurement itself. Here, we describe both approaches.

For the reference measurement, we used a doughnut-shaped illumination profile but the calibration of the orbital scan can be done with a Gaussian profile as well. We focused on a single GNP and zoomed in, ultimately leading to a situation in which the galvo is not scanning anymore, but an image of the GNP can be seen on the array detector. Then, using the LabVIEW software, we started a circular scan around that position, keeping the orbit period (1.92 s) and induced radius 100 nm) the same as for the ISM-FLUX measurements. We measured for about 19 seconds and skipped the first couple of seconds in the analysis to remove galvo artifacts at the start of the scan. The laser power was chosen to have a photon count rate of about 7 MHz overall detector elements combined, orders of magnitude higher than the dark count rate but low enough not to induce saturation effects. As a result of the descanned detection mode, the movement of the galvo is reflected in the position of the particle on the array detector. To find the orbit parameters, we first summed, for each position in the orbit, the photon counts of 100 orbits. Then, we localized the particle for each position in the orbit with the maximum likelihood estimator, resulting in a circle of 32 points. We repeated this procedure for the whole data set, leading to a set of circles, from which the radius, starting angle, and orientation can be measured as explained in the protocol of (Fig. S8). Note that, due to inertia, the actual orbit radius and starting angle may significantly differ from the imposed values, (Fig. S13).

All orbit parameters can also be directly retrieved from the SM data (Fig. S9). Since the detector is placed in descanned mode, moving the illumination beam leads to an identical apparent movement of the detector in the sample space and thus leads to a corresponding opposite shift in the image of a molecule. In other words, localizing each on-event for every angle of the TCP individually results in 32 distinct localizations, each exhibiting a relative shift with respect to the other 31 localizations. From the resulting pattern, one can derive all orbit parameters, *i*.*e*., the diameter, the starting angle, and the direction of rotation. Given the low number of photon counts per position in the TCP, we modified this concept by binning the photon counts in 4 quadrants of 8 consecutive points of the TCP, resulting in 4 sets of localizations rather than 32. We performed the localization analysis for different *L* values and calculated for which *L* value the relative shift between the four sets is the smallest. The resulting value was in good agreement with the true *L* value that we measured in a reference measurement, (Fig. S8).

### G. Data analysis

All data analysis was done in Python. A scheme of the analysis protocol is shown in (Fig. S14) and (Fig. S15). For the origami measurements, we started by calculating a time trace, (Fig. S14) (a), obtained by summing all photons in all detector channels (excluding the two that have high dark-noise, (Fig. S7)) for all positions on the orbit. Then, we calculated a histogram, which was fitted with a sum of two Gaussian distributions as a fit function. The count value corresponding to the first peak was used as a threshold to distinguish the signal from the background. Each time the photon counts exceeded the threshold, a new event started. The event ended when the intensity trace fell again below the threshold. For all orbits within an event, photon counts observed in the same detector element for the same position in the orbit were summed, resulting in a reduced event data set *n*_*ij*_, with *I* ∈{1, 2, …, *N*_*c*_} and *j* ∈{1, 2, …, *N*_*d*_}, with *N*_*c*_ = 32 and *N*_*d*_ = 23. The events were further filtered, (Fig. S14) (b), based on the total number of photons, the duration, and the standard deviation *σ*_measured_ of the count fluctuations within an event. Given a high background count rate stemming from a combination of out-of-focus fluorescence and detector dark counts, a threshold of at least 2000 photons per event was enforced. Furthermore, only events with a duration of at least 5 orbits were considered. Lastly, we calculated for each event the standard deviation of the photon counts over the orbits and discarded events for which this standard deviation was more than 2.5 times the expected value for a Poisson process, *i*.*e*., 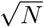. This filter reduces the chance of including events characterized by pronounced blinking or the presence of a secondary fluorophore bound to a nearby docking site. Each event that passed all filters was split into three equally long segments, (Fig. S14) (c), resulting in three independent localizations. If the three localizations were closer to each other than a user-chosen threshold, *i*.*e*., *σ < σ*_*T H*_, the event was accepted; otherwise, the event was discarded. Here,),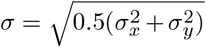 with *σ*_*x*_ and *σ*_*y*_ the standard deviation of the *x* and *y* locations, respectively.

For the measurements with the fiducial markers, no event detection was needed. Instead, we moved the marker every second (Fig. 3) (a-e) or half a second, panel (f), with the piezo-electric stage to a pre-programmed position while continuously performing the orbital scan and detecting the signal with the array detector. The resulting trace was binned several times with increasing bin lengths to increase the photon counts. For ((Fig. 3)) (a-c, f), each of the 17 positions was probed 50-100 times, and the standard deviation in *x* and *y* was used to calculate the localization uncertainty. For (Fig. 3) (d), we probed each location at least 39 times, and we calculated the covariance matrix of the localizations and used the eigenvectors and the square root of the eigenvalues of the matrix to find the orientation and size of the principal components of the spread, respectively.

For the measurements with the fixed fluorophores, the intensity trace was binned several times with increasing bin lengths to increase the photon counts. For a given bin length, each set of three consecutive bins was analyzed as follows: first, the fluorophore position was calculated for each of the three events. The standard deviation in (x,y) was used as an estimate for the localization uncertainty for that set. The final localization uncertainty for a given *N* was calculated as the mean ± standard error over all sets. Similarly, *N* was calculated as the mean ± standard error over all sets. Depending on the bin length, between 22 and 366 events of 75-1300 photons (fluorophore position 1) and between 50 and 2500 events of 43-2200 photons (fluorophore position 2) were analyzed.

Localizations were calculated with a brute-force maximum likelihood approach, similar to (10). Each event consisted of either 32×25 or 32×23 photon count values, with and without hot detector elements, respectively, and an equally large set of MDFs. For the GNP and the 20 nm origami, we use the experimental MDFs; for the 40 nm origami the theoretical MDFs. We analyzed all events in a data set assuming the same SBR, set to 3 for the fiducial marker, 3.13 for the 40 nm origami, and 3.3 for the 20 nm origami, in all cases assuming the background to be uniformly distributed over all positions in the orbit and all detector elements.

## Data and code availability

An exemplary ISM-FLUX data file and a Jupyter Notebook with analysis instructions to replicate the results have been deposited on Zenodo (36).

## ACKNOWLEDGEMENTS

The authors thank L. Masullo, J. Stein, and F. Balzarotti for advice and fruitful discussions and G. Garrè for helping with the analysis. This research was supported by the European Research Council, BrightEyes No. 818699 (E.S., S.P., M.O.H., G.V.) and the European Union’s Horizon 2020 research and innovation program under the Marie Sklodowska-Curie Grant Agreement No. 890923 (SMSPAD) (E.S.).

## AUTHOR CONTRIBUTIONS

G.V. and E.S. conceived the idea. E.S. and S.P. built the setup and performed the experiments. G.V., E.S., and S.P. analyzed the data. M.O.H. prepared the fixed molecule sample. A.Z. wrote the code to extract the detector orientation, rotation, and magnification from an ISM data set. All authors contributed to the writing.

## Supplementary Note 1: ISM-FLUX theory

Consider a system in which a fluorophore, i.e., a point-emitter, is sequentially illuminated with *K* illumination patterns, e.g. by moving a doughnut beam to different positions. The number of photons detected with a single-element detector can be assumed to be Poisson distributed with *λ*_*i*_ the expected number of photons under illumination *i* (*I* ∈ {1, 2, …, *K*}). Thus, the probability of observing **n** = (*n*_1_, *n*_2_, …, *n*_*K*_) photons is

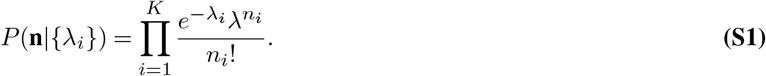

However, this is the probability of having **n** photons. What we are interested in is the probability *P* (**n**|*N*), with *N* = ∑_*i*_ *n*_*i*_. Using Bayes’ theorem, we have

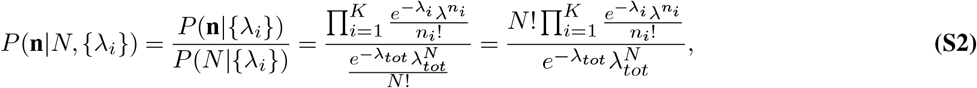

with *λ*_*tot*_ = ∑*λ*_*i*_.

We can write *λ*_*i*_ = *Np*_*i*_, with *N* the total number of expected detected photons, *p*_*i*_ the probability that if a photon is detected, it is detected under illumination *i*, and ∑_*i*_ *p*_*i*_ = 1. Then,

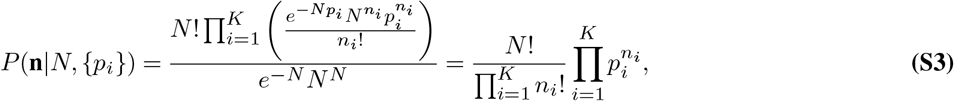

which is a multinomial distribution.

We now turn to the ISM-FLUX case with an array detector. For ISM-FLUX, we can define a probability *p*_*ij*_ that if a photon detection event happens, it happens under illumination *i* in detector-element *j*. We can write the dependence of *p*_*ij*_(**r**_*E*_) on the emitter position **r**_*E*_. Assume a (doughnut) illumination intensity profile *h*(**r**−**r**_*i*_) that moves to *K* different positions **r**_*i*_. Assume further that the expected number of emitted photons is proportional to the illumination intensity. Then, the fraction of signal photons emitted under illumination *i* is:

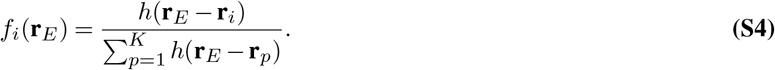

The fluorescence emitted by the single emitter follows a spatial distribution *h*_2_(**r**−**r**_*E*_) equal to the emission PSF centered at position **r**_*E*_. We have,

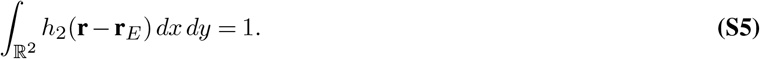

The probability *g*_*ij*_(**r**_*E*_) that a photon emitted under illumination *i* is detected by detector element *j* is

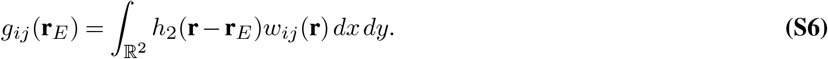

Here, *w*(**r**) is a 2D window function describing the detector element. Note that, since the detector is placed in descanned mode, the detector *moves* when the excitation beam moves, hence the dependence on both the indices *i* and *j*:

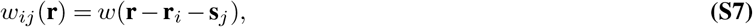

with **s**_*j*_ the position of detector element *j* within the array. Note that the detector has a finite number of elements and thus a finite size, hence not all emitted photons will be detected:

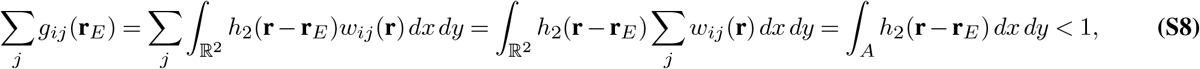

with *A* the overall area of all detector elements.

If a photon is observed, the probability it happened under illumination *i* in detector element *j* is:

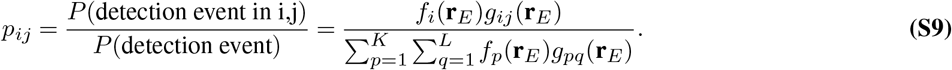

Plugging in S4 yields

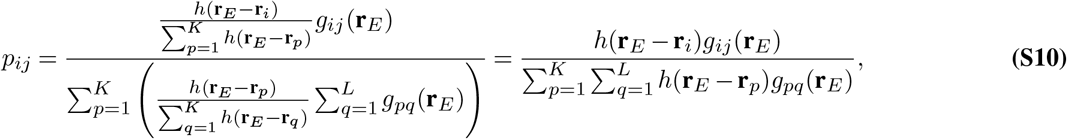

where, using Eq. S6, the numerator can be written as

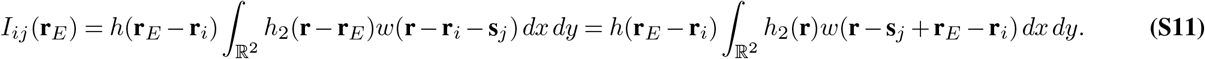

For **r**_*i*_ = 0, we find the equation for the signal in detector element *j* for a point-source at position **r**_*E*_ and detector element at position **s**_*j*_. Experimentally, one can measure *I*_0*j*_(**r**_*E*_) by moving a point source in 2D with a piezoelectric stage and measuring the resulting signal for each position and in each detector element or, equivalently, by scanning the point source with the galvanometric scanners.

For **r**_*i*_*/=*0, we have *I*_*ij*_(**r**_*E*_) = *I*_0*j*_(**r**_*E*_ −**r**_*i*_). Thus, when the TCP is known, one can simply shift the resulting point-source images to the right positions to get the full set of *I*_*ij*_(**r**_*E*_). One can find the denominator in Eq. S10 by summing all images. For the ISM-FLUX setup in this work, we have *K* = 25 detector elements and *L* = 32 positions.

In the presence of background, Eq. S10 becomes:

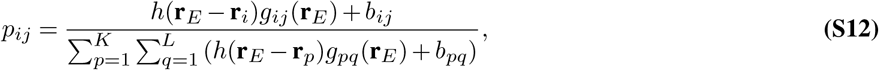

with *b*_*ij*_ the probability that a count is a background count in element *j* under illumination *i*. With a SPAD array detector, the main contributions to the background counts are the out-of-focus fluorescence and the dark counts. We assume that both contributions are equal for each detector element and each illumination, thus *b*_*ij*_ = *b*:

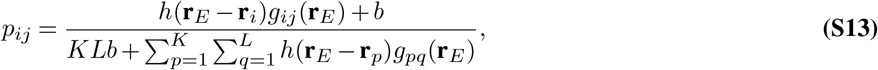

Define the signal-to-background ratio as:

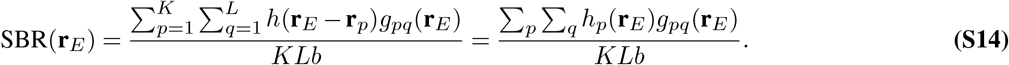

where we introduced a shorter notation for the double sum. Then,

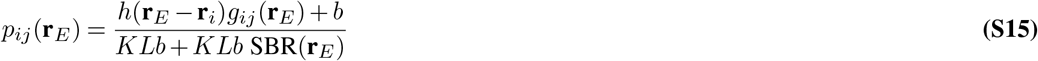

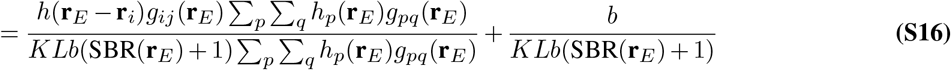

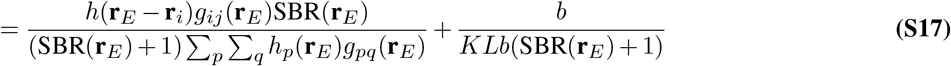

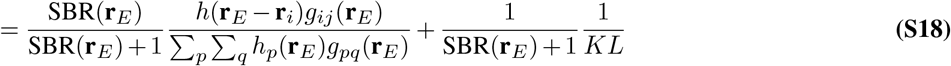

Assuming no prior information, we can express a likelihood function ℒ as

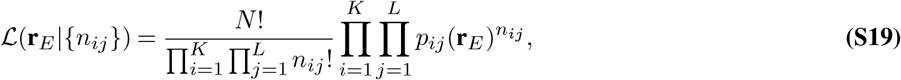

with *p*_*ij*_ from Eq. S18.

The log-likelihood function, dropping constant terms, is

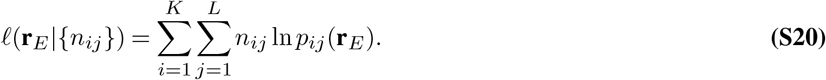

The maximum likelihood estimation for the emitter position is

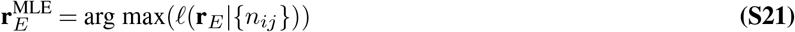

## Supplementary Note 2: Bias of the maximum-likelihood estimator in ISM-FLUX

The maximum-likelihood estimator (MLE) is not inherently unbiased. Especially in the case of small samples, i.e. low photon counts, there may be a difference between the expected value of the emitter position and the ground truth value. To understand the bias of the MLE for ISM-FLUX, we simulated ISM-FLUX measurements for different emitter positions and different photon counts and compared the position retrieved from the MLE with the ground truth. We consecutively positioned the emitter at various points along a line passing through the TCP center. We used the simulated MDFs, Fig. S11, to draw photon counts from a multinomial distribution with relative probabilities equal to the MDF values at the emitter position. We assumed 0 background. Then, we used the MLE to retrieve the emitter position and stored the difference in (*x, y*) between the ground truth and the retrieved position. We repeated this simulation 1000× for each emitter position and each photon count number.

The result, Fig. S1, shows that the MLE is always unbiased in the TCP center and on the TCP circle, regardless of the number of photon counts. For other positions within the TCP, as well as outside the TCP, we find a position- and photon-count-dependent bias. For high photon counts, i.e. *N >* 1000, the bias is far below 1 nm in both directions for all positions within the detector FOV and the MLE can be considered unbiased. For lower photon counts, the bias increases in both directions but remains below 10 nm for the whole FOV for *N* ≥ 30. Note that for most positions on the horizontal line, the bias in the *x* (radial) direction is higher than in the *y* (tangential) direction.

**Fig. S1.**
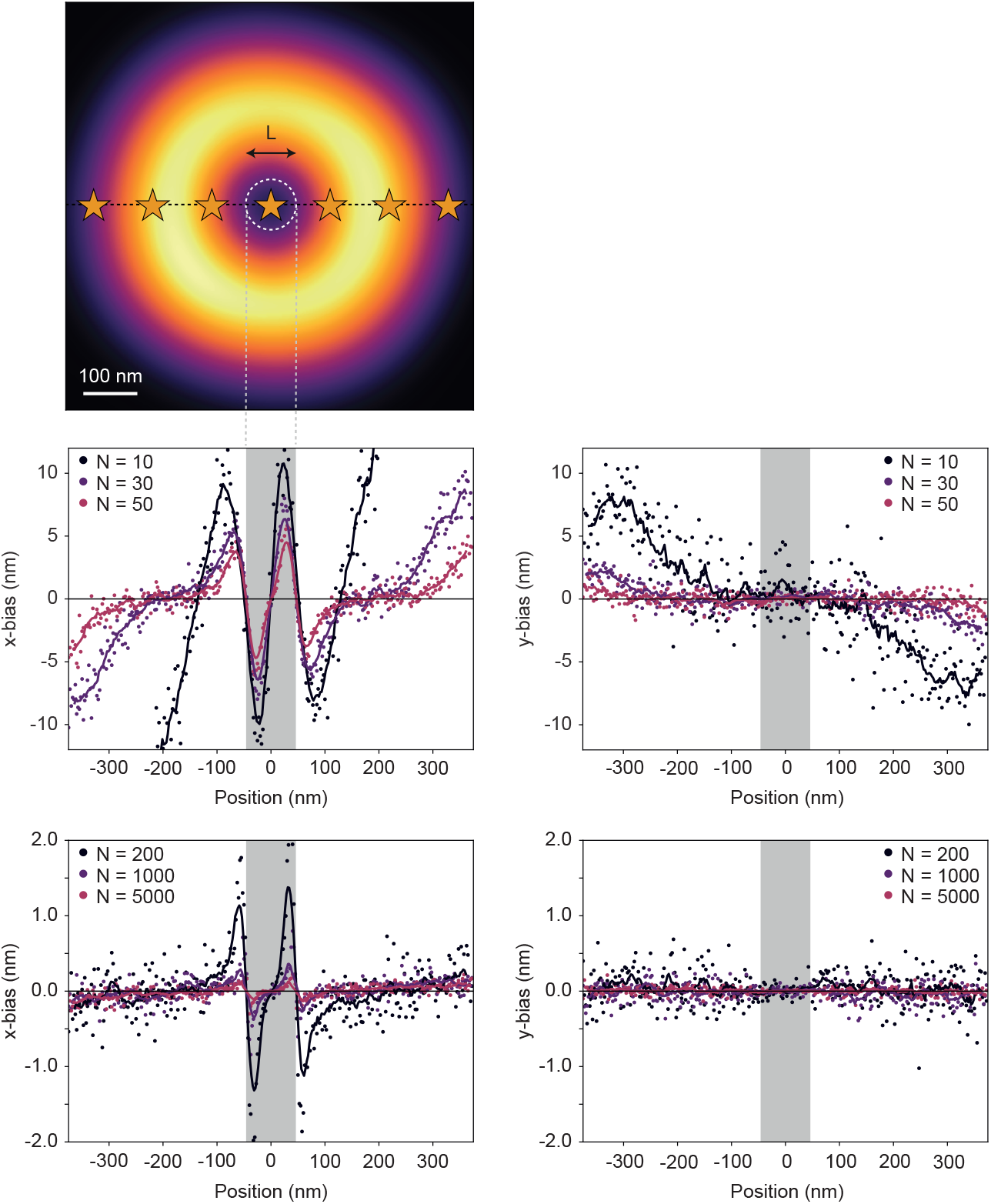
Bias of the MLE in the *x* and *y* directions for different photon counts (*N*) and various emitter positions. *L* = 90 nm. The emitter is moved to 300 positions in 2.5 nm steps along a horizontal line passing through the TCP center (indicated by the dotted black line in the intensity plot). For each position, the bias in both directions is plotted. The scatter plots display the raw data, while the line plots show the corresponding moving averages with a window size of 10.

## Supplementary Note 3: Theory of ISM-FLUX self-calibration

Consider a molecule at position **r**_*E*_ and two opposite points on the TCP, located at **r**_1_ and **r**_16_. When the laser beam is at position **r**_1_ or **r**_16_, the detector center is also at **r**_1_ or **r**_16_, respectively. Thus, in the detector frame-of-reference and assuming no (shot) noise, the image of the molecule moves from being centered around **r**_*E*_− **r**_1_ to **r**_*E*_− **r**_16_. The shift of the molecule **L** as seen by the detector is:

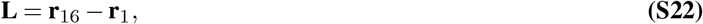

with |**L**| = *L*, the diameter of the TCP. Thus, *L* can be directly derived from two micro-images taken at opposite angles of the TCP.

However, **r**_1_ and **r**_16_ are experimental localizations subject to a non-zero localization uncertainty. As a more robust approach, we propose to localize events for different points on the TCP assuming a TCP diameter *£*. In other words, we shift the two emitter positions in the detector reference frame with ±***£****/*2, with ***£*** a vector on the line connecting the two points on the TCP and |***£***| = *£*:

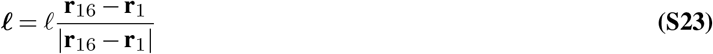

The two emitter positions are then

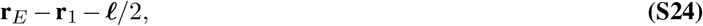

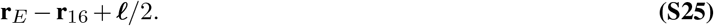

The difference Δ**r** between the two positions is

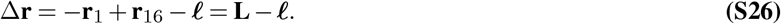

For ***£*** = **L**, Δ**r** = 0.

In reality, **r**_1_ and **r**_16_ are localizations with uncertainty *σ*^2^. Assuming the localizations can be drawn from a Gaussian distribution with standard deviation *σ*, the mean squared distance ⟨*d*^2^⟩ between the two positions and their mean is

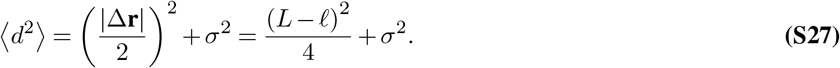

Thus, ⟨*d*^2^⟩ reaches a minimum where *£* = *L* and the function value at this point corresponds to the experimental localization uncertainty.

Since the photon counts in a single-molecule on-event are rather low, we split the data from each event into 4 circle segments of 8 consecutive points in the orbit. Then, we approximate ⟨*d*^2^⟩ as the mean squared displacement between the four localizations and their mean. As a metric for the overall localization spread of all events combined, we take the median value of all ⟨*d*^2^⟩ values. We repeat this calculation for a set of *£* values and we fit the resulting curve with a second-order polynomial. The *£* value that minimizes the localization spread corresponds to the true *L* value. The protocol is illustrated in Fig. S9.

**Fig. S2.**
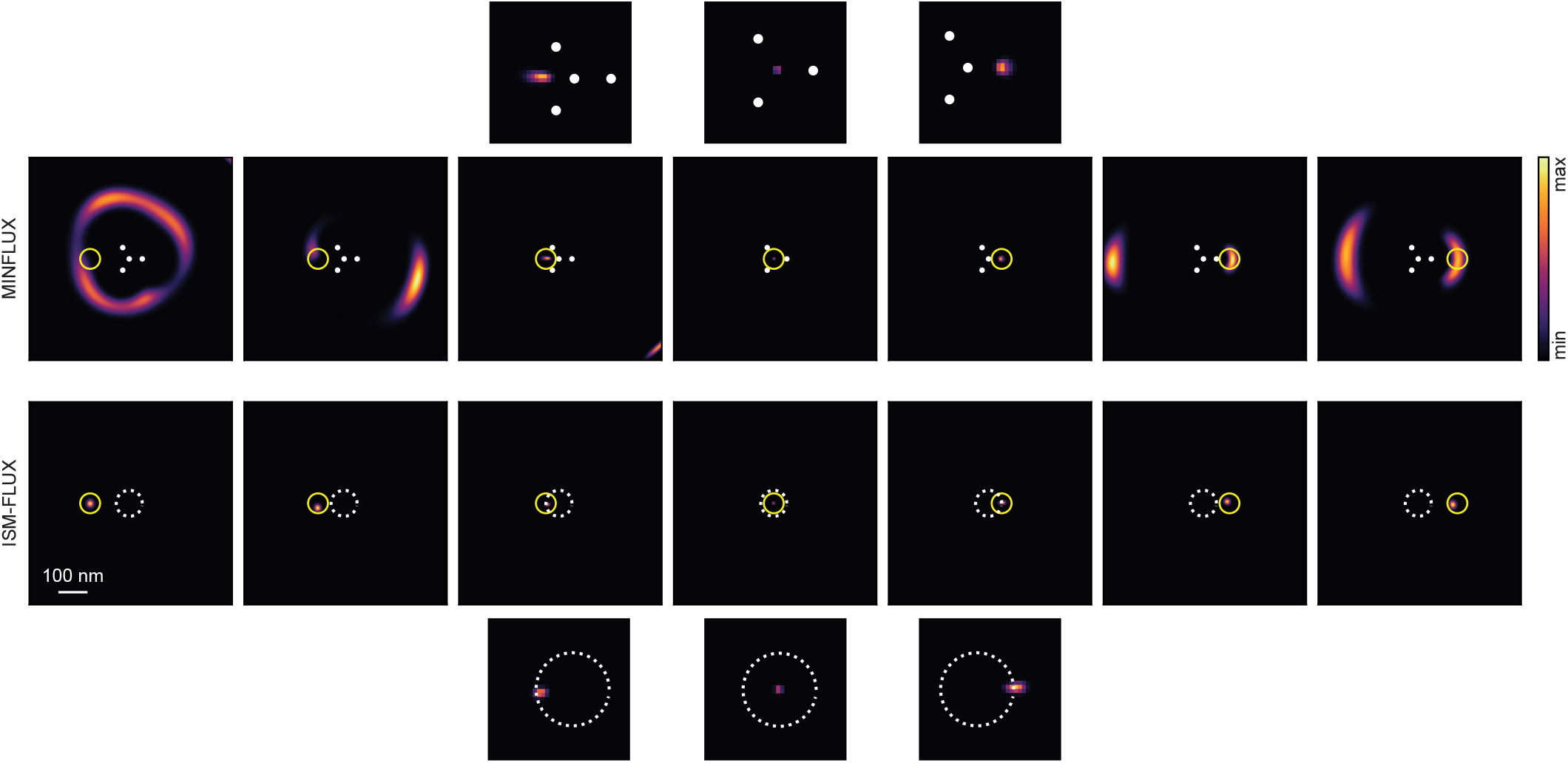
Likelihood maps for simulated MINFLUX and ISM-FLUX measurements. For MINFLUX, we considered a TCP, indicated in white, consisting of 3 evenly spaced points on a circle with a diameter *L* = 90 nm and one point in the center of this circle. For ISM-FLUX, the TCP consists of 32 points on a circle with a diameter of 90 nm, without the central point. We placed the emitter at 7 different positions in steps of 45 nm, indicated by the yellow circles, and we calculated the MLE maps for *N* = 200 photons. For visualization purposes, some TCP points are not shown. The top and bottom row images show zoomed-in MLE maps of the three emitter positions closest to the TCP center.

**Fig. S3.**
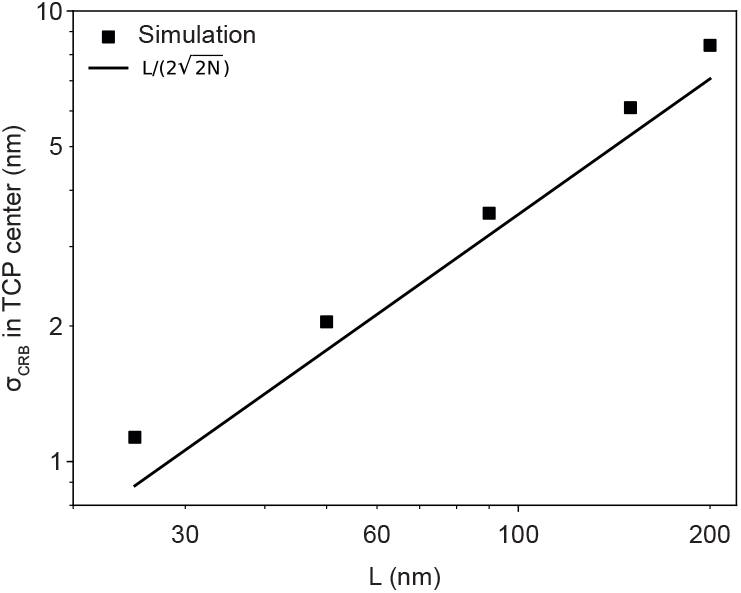
Simulated *σ*_CRB_ in the TCP center as a function of *L* for 100 photons. The full line corresponds to the theoretical MINFLUX localization uncertainty for 100 photons.

**Fig. S4.**
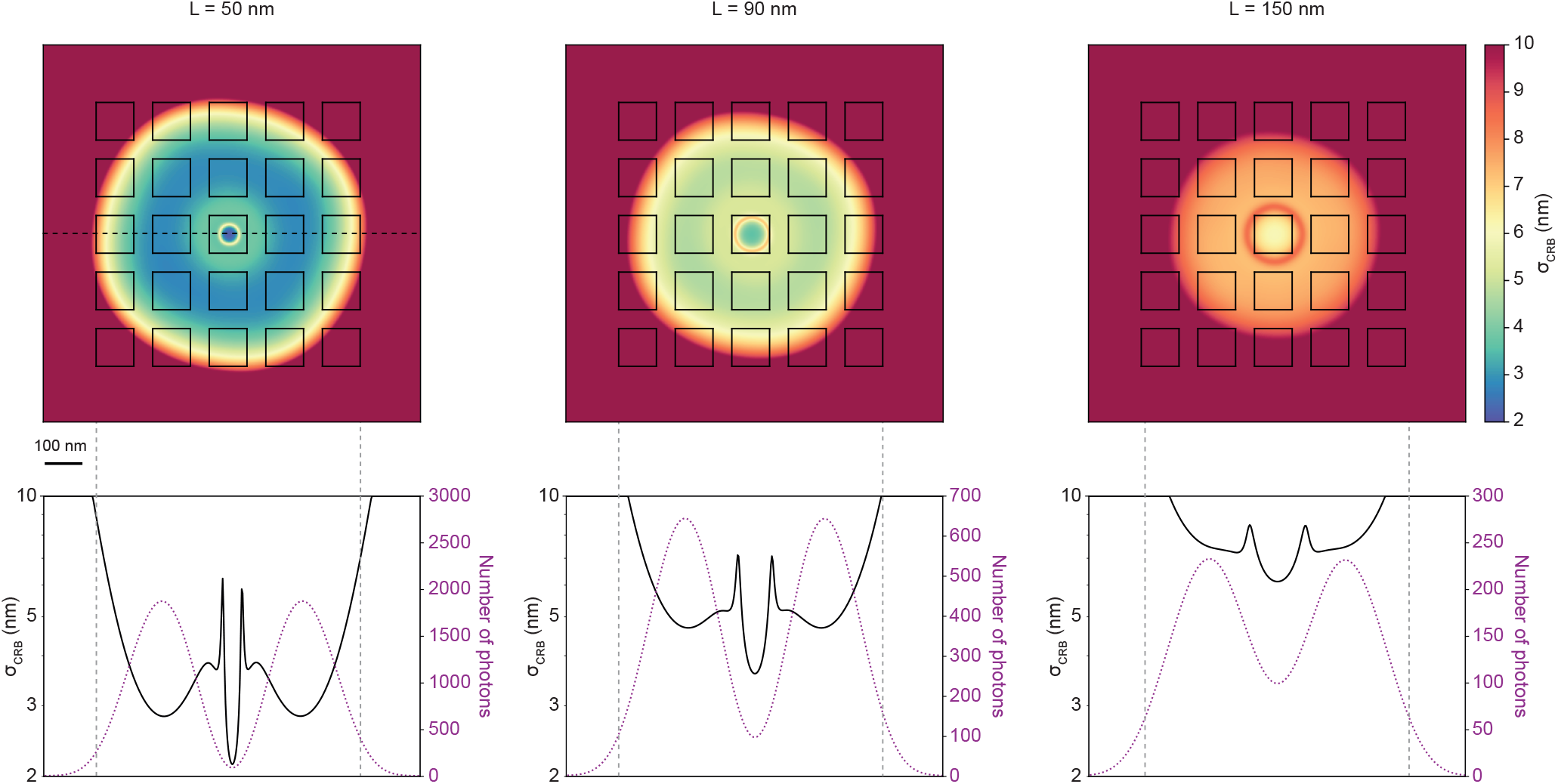
CRB for a molecule emitting *N* = 100 photons in the TCP center. *N* is rescaled for the other emitter positions, assuming *N* (**r**) is proportional with 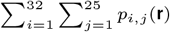.

**Fig. S5.**
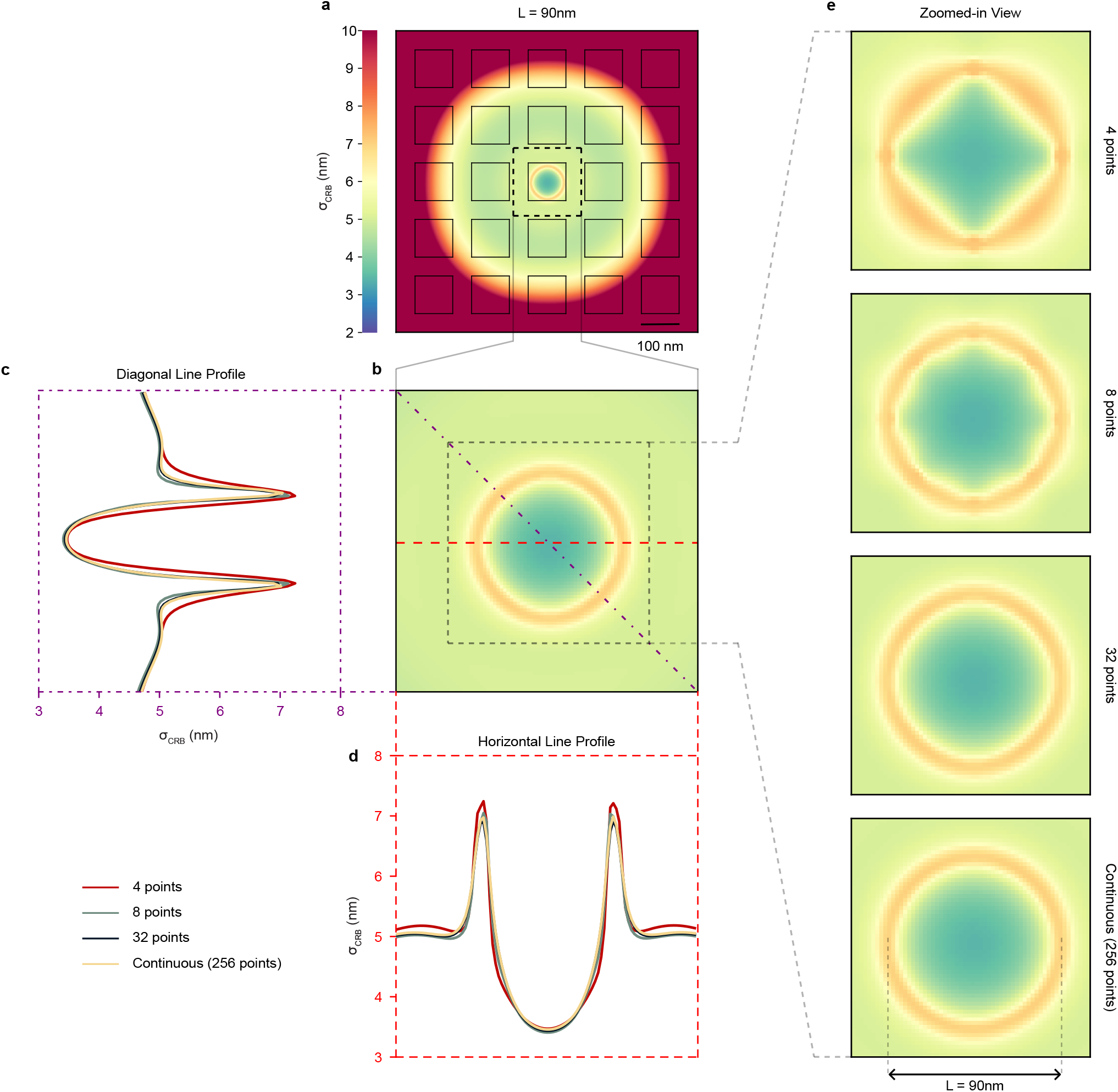
(a) CRB for a molecule emitting *N* = 100 photons in the TCP center. *N* is rescaled for the other emitter positions. *L* = 90 nm. (b) Zoomed-in section of the center of the CRB map. (c) *σ*_CRB_ along the diagonal line in (b) and comparison with other circular TCPs discretized with a different number of points. (d) Same as (c) for the horizontal line profile of *σ*_CRB_, indicated by the red line in (b). (e) *σ*_CRB_ maps for circular TCPs discretized with a different number of points.

**Fig. S6.**
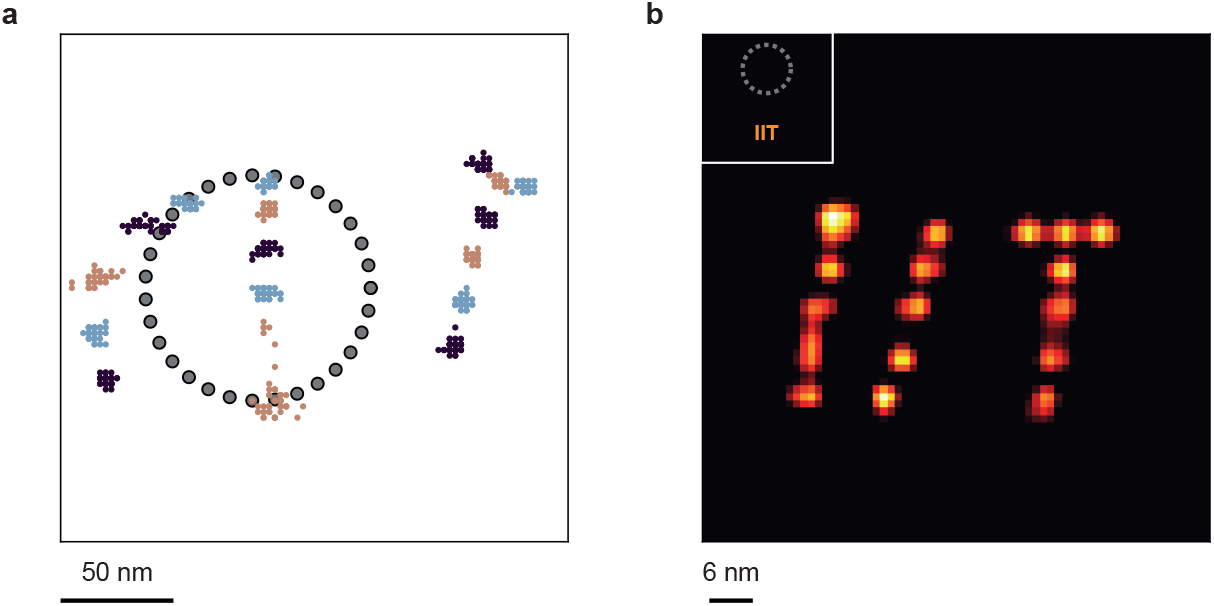
GNP results for simulated MDFs. For (b), the GNP from the time was outside the TCP, as indicated in the inset.

**Fig. S7.**
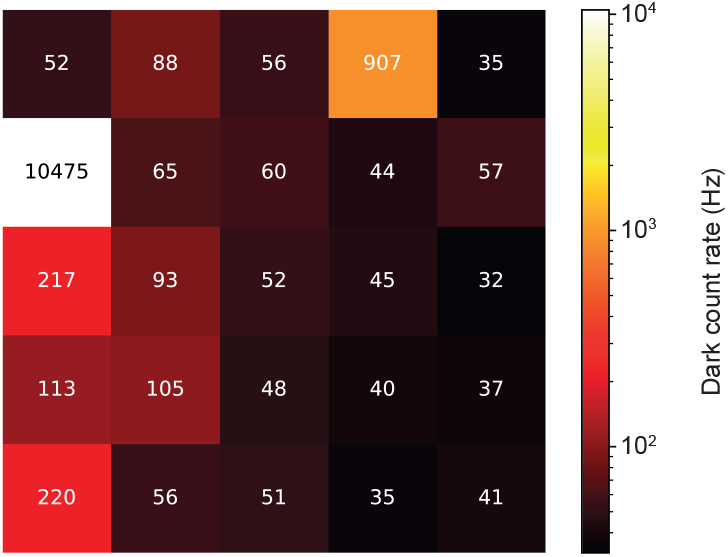
Dark count rate for the 25 most central detector elements of the 7×7 array. Pixel 3 (fourth pixel of the first row), and pixel 5 (first pixel of the second row) are hot elements and were – unless mentioned otherwise – excluded from the data analysis. Dark count rates for each channel were obtained by covering the detector and fitting to a line the photon counts accumulated over a two-second period.

**Fig. S8.**
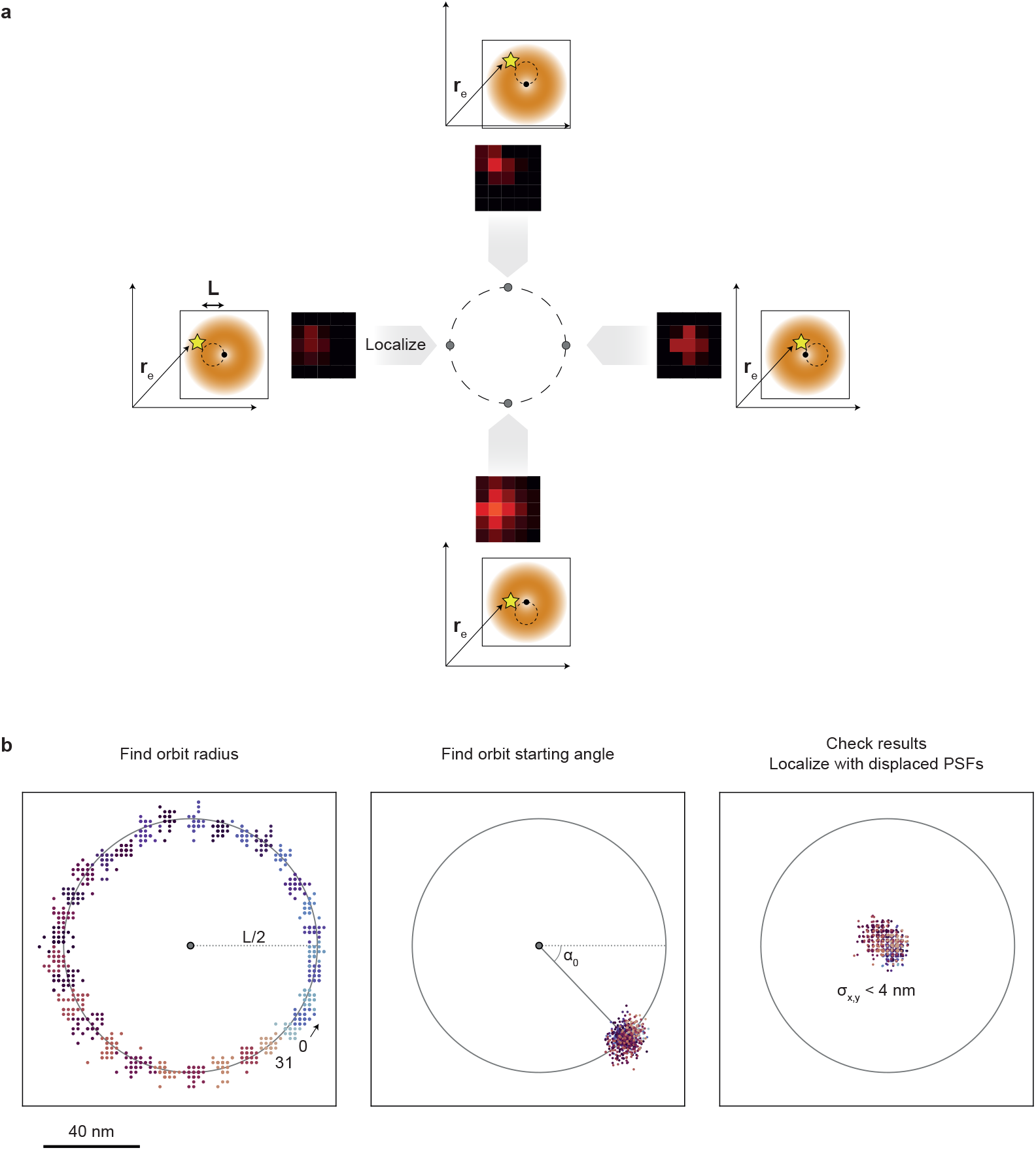
Measurement of the circular orbit parameters from a reference measurement. (a) A fiducial marker is imaged while scanning the sample with a circular motion. Each position on the circle leads to a different micro-image, both in terms of the center of mass as well as the number of photons. The marker is localized in each micro-image individually using the 5×5 PSFs. Note that in this case only the distribution of the scattered photons on the detector is used, not the intensity differences between different images. Hence, also a Gaussian beam can be used for calibrating the galvanometric scanners. (b) Analyzing the 32 micro-images using an MLE leads to a distribution of localizations on a circle from which the center is estimated by taking the mean coordinates and the radius is estimated as the mean distance from the center. Mean and standard deviation over 30 circles yields *L* = (105.2 ± 0.9) nm. Next, all points are rotated back an integer number of 1*/*32 ∗ 2*π* and from the mean resulting coordinates, the starting angle of the orbit is estimated. Having both the orbit radius and starting angle, a new series of PSFs can be calculated: one set of 25 PSFs for each of the 32 positions on the circle, calculated by displacing each set of 25 PSFs according to its position in the orbit. The resulting set of 32×25 PSFs is used to analyze ISM-FLUX data, i.e. data from a full orbit or a sum over several consecutive orbits will result in a single localization, in which both the distributions of the photons on the detector as well as the change in photon flux during the orbit is taken into account.

**Fig. S9.**
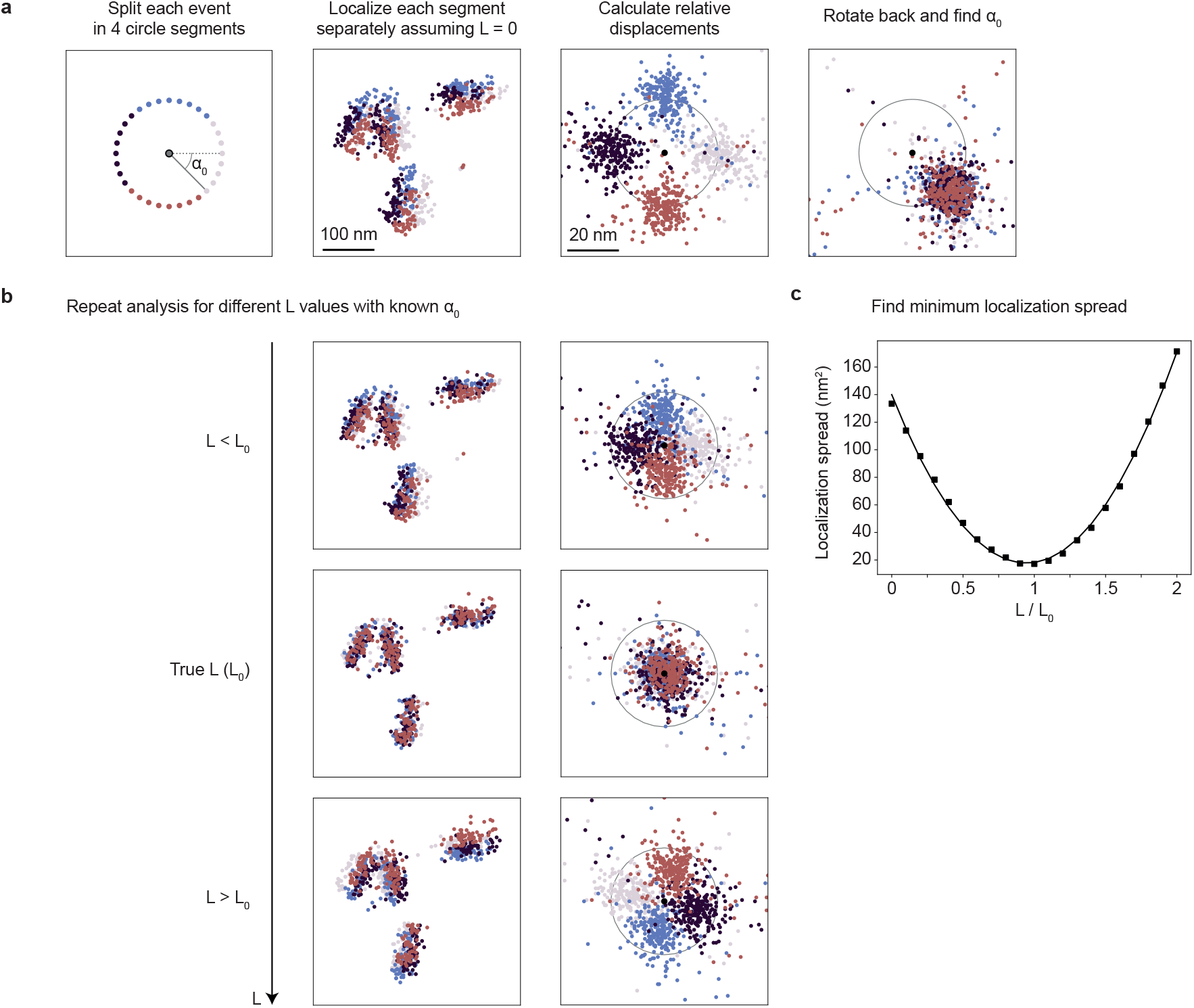
ISM-FLUX is self-calibrating: measurement of the orbit parameters from the SM data. (a) Extracting the starting angle of the orbit *α*_0_. Each blinking event is split into four segments, indicated by the different colors. The MLE is applied to each segment separately assuming *L* = 0. The result is four sets of segment-localizations that are displaced with respect to each other. Calculating for each event the relative displacement between the segments (i.e., the displacement between the localization of each segment and the mean of the four segments) results in four clusters on a circle. By rotating each segment *s* back by an amount equal to −*π/*4 − *sπ/*2 with *s* ∈ {0, 1, 2, 3}, all clusters overlap and the starting angle *α*_0_ can be measured. We calculated the median (x,y) position of each cluster and estimated *α*_0_ as the mean ± standard deviation, which is *α*_0_ = −0.81 ± 0.02, in good agreement with the value of *α*_0_ = −0.812 ± 0.008 found in a reference measurement, Fig. S8. (b) The orbit diameter is found by repeating the analysis assuming different *L* values. For each *L* value, the localization spread, as defined in Sup. Note 3, is calculated from the relative displacements. (c) The localization spread as a function of *L* (scatter plot) is fitted with a second-order polynomial curve (line plot). The fit, *y* = 137*x*^2^ 259*x* + 140 has a minimum around *L/L*_0_ = 0.95, with *L*_0_ = 61.4 nm, the diameter found in the reference measurement.

**Fig. S10.**
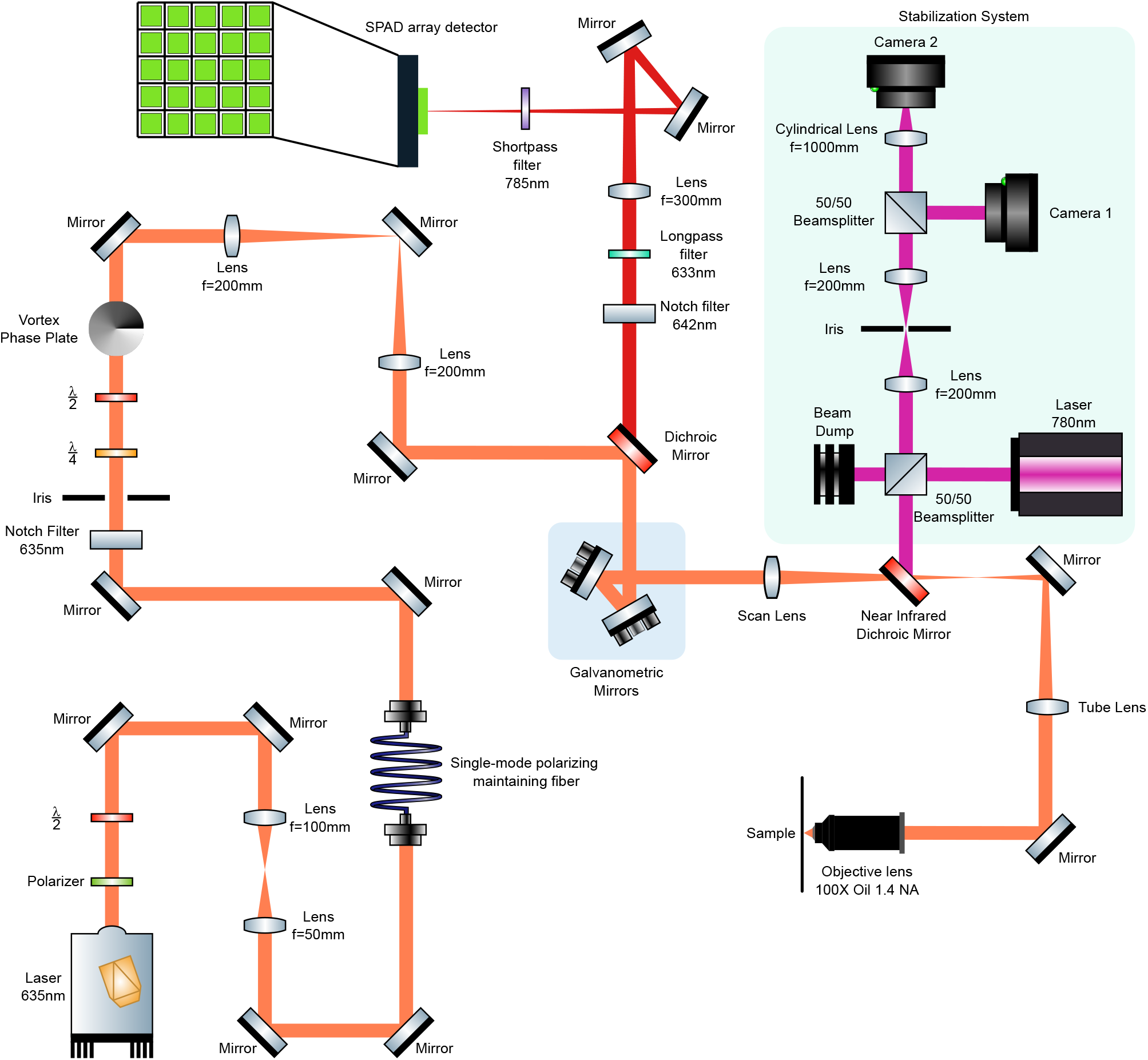
Detailed optical setup 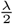 and 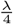 stands for half-wave and quarter-wave plate. Near infrared dichroic mirror used in this system is Shortpass 750nm. The optical setup highlighted in green is the stabilization system used to keep the sample in focus during the imaging session.

**Fig. S11.**
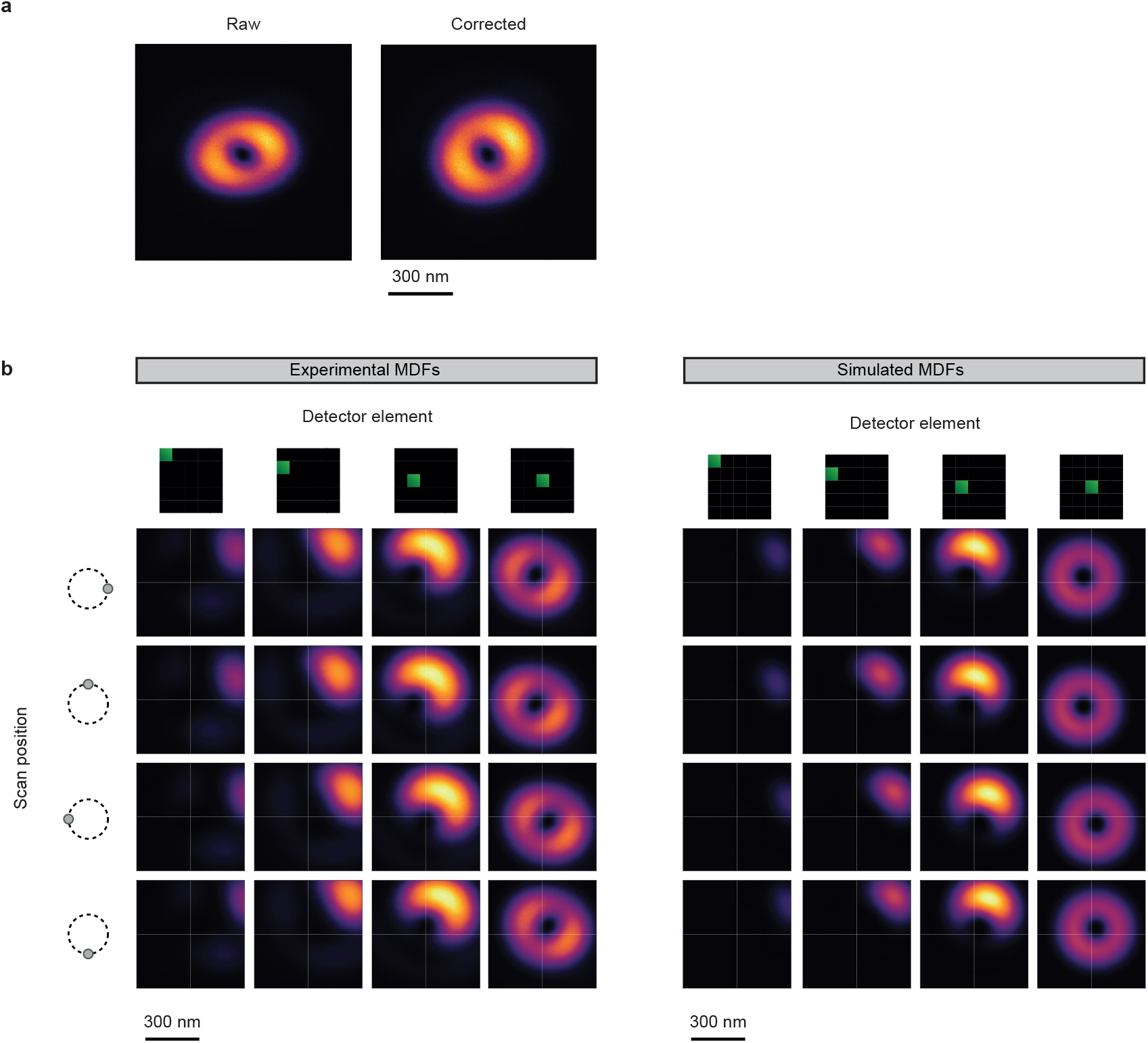
(a) Experimental MDF for the central detector element, obtained by scanning a gold nanoparticle. Scan settings… Galvanometric limitations lead to an apparent astigmatic image, corrected in post-processing. (b) Subset of (a) experimental and (b) simulated MDFs. The full MDF data set is a collection 25 x 32 images, a combination of 25 detector elements and 32 scan positions. For simplicity, we show 4 positions and 4 detector elements. The experimental MDFs are stretch-corrected and smoothed in post-processing.

**Fig. S12.**
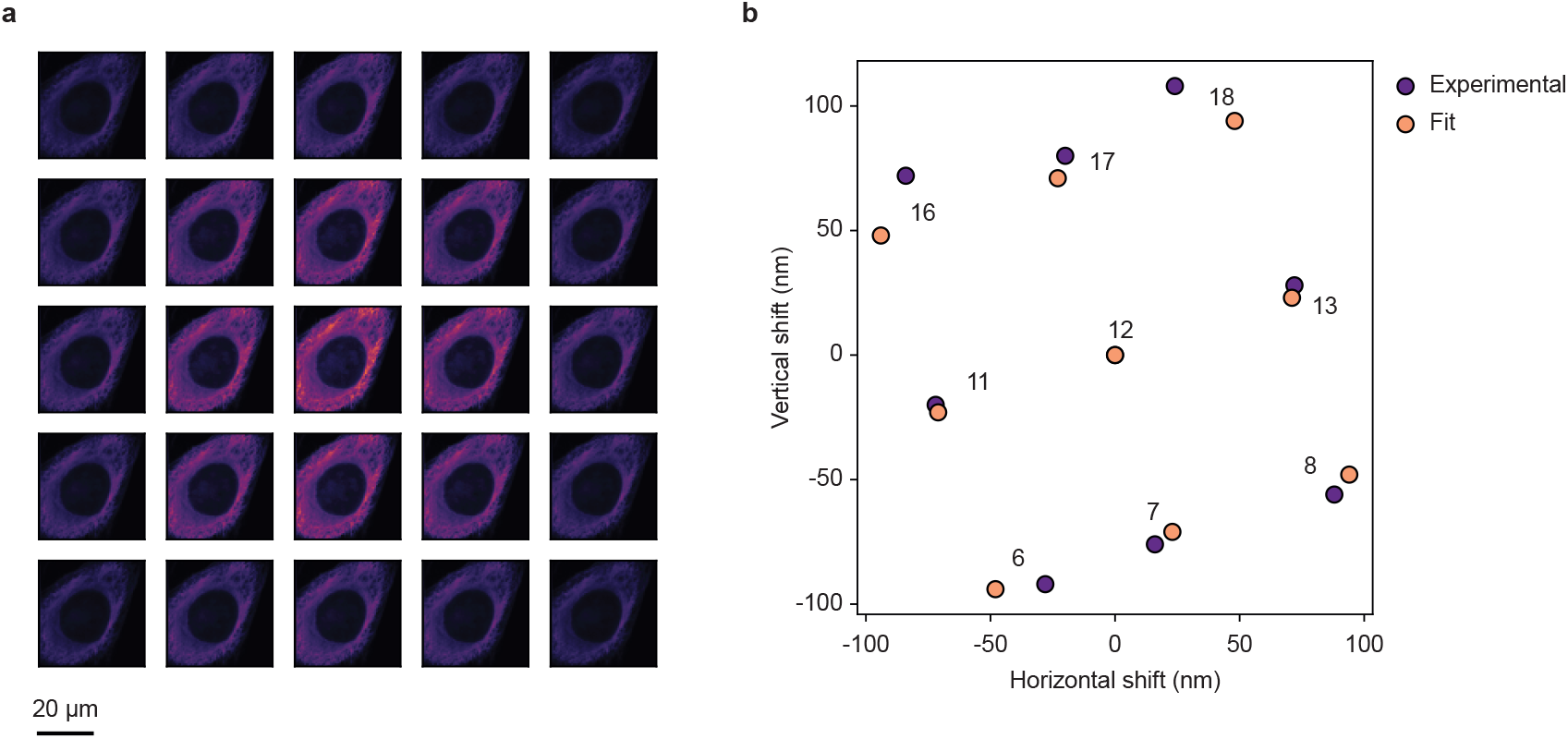
Extracting the detector orientation, rotation, and system magnification from the shift vectors of an ISM data set. (a) Image of a fixed HeLa cell with *α*-tubulin staining. The images of the 5×5 detector elements are shown. (b) From the shift vectors of the 3×3 detector elements closest to the detector center, the detector orientation, rotation, and system magnification were estimated.

**Fig. S13.**
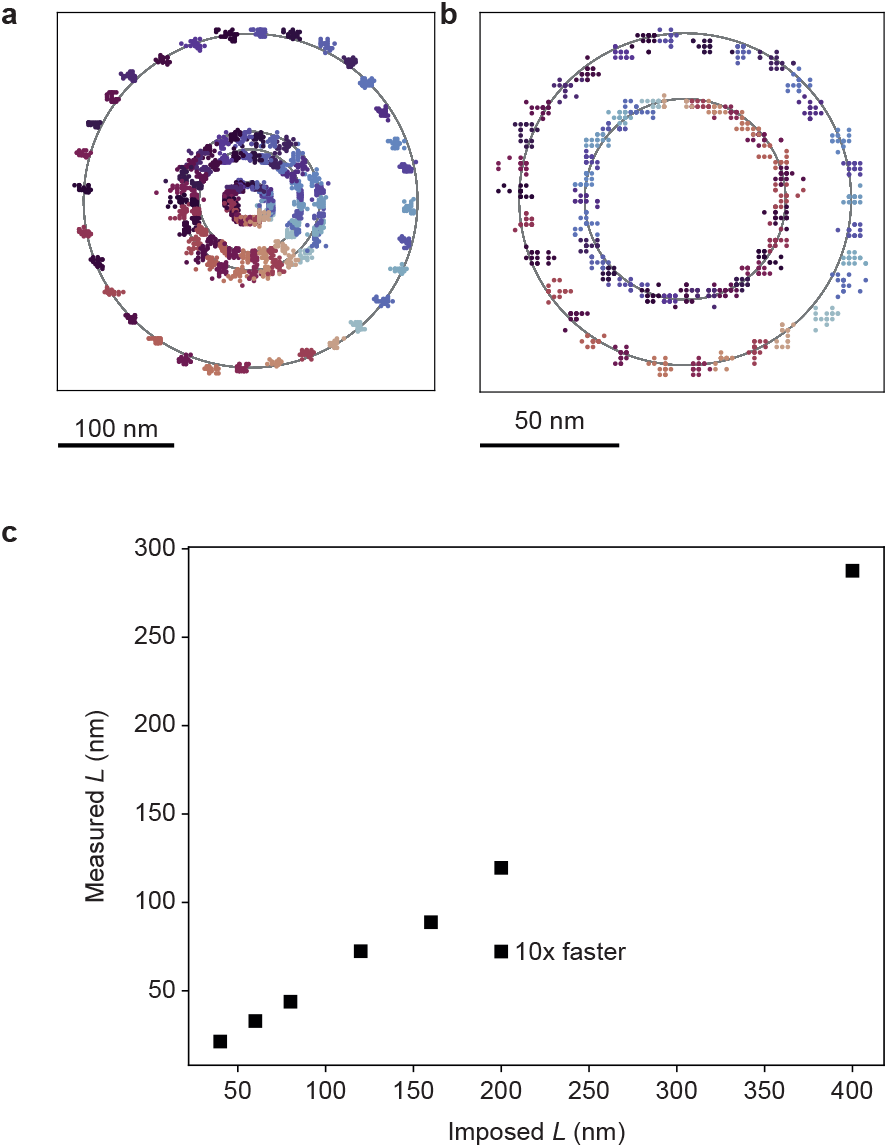
(a) 32 positions of the TCP measured for different *L* values. From smallest to largest TCP: *L* = 60 nm, 160 nm, 200 nm, 400 nm. Orbit direction counterclockwise, from blue to red. (b) The same imposed *L* of 100 nm results in a different observed *L* and starting angle, depending on the scan speed. The smaller circle corresponds to a 10x faster scan (192 *µ*s per circle) than the larger circle (c) Imposed *L* vs. measured *L*. Except for the indicated data point, all orbit times were 1.92 ms per circle.

**Fig. S14.**
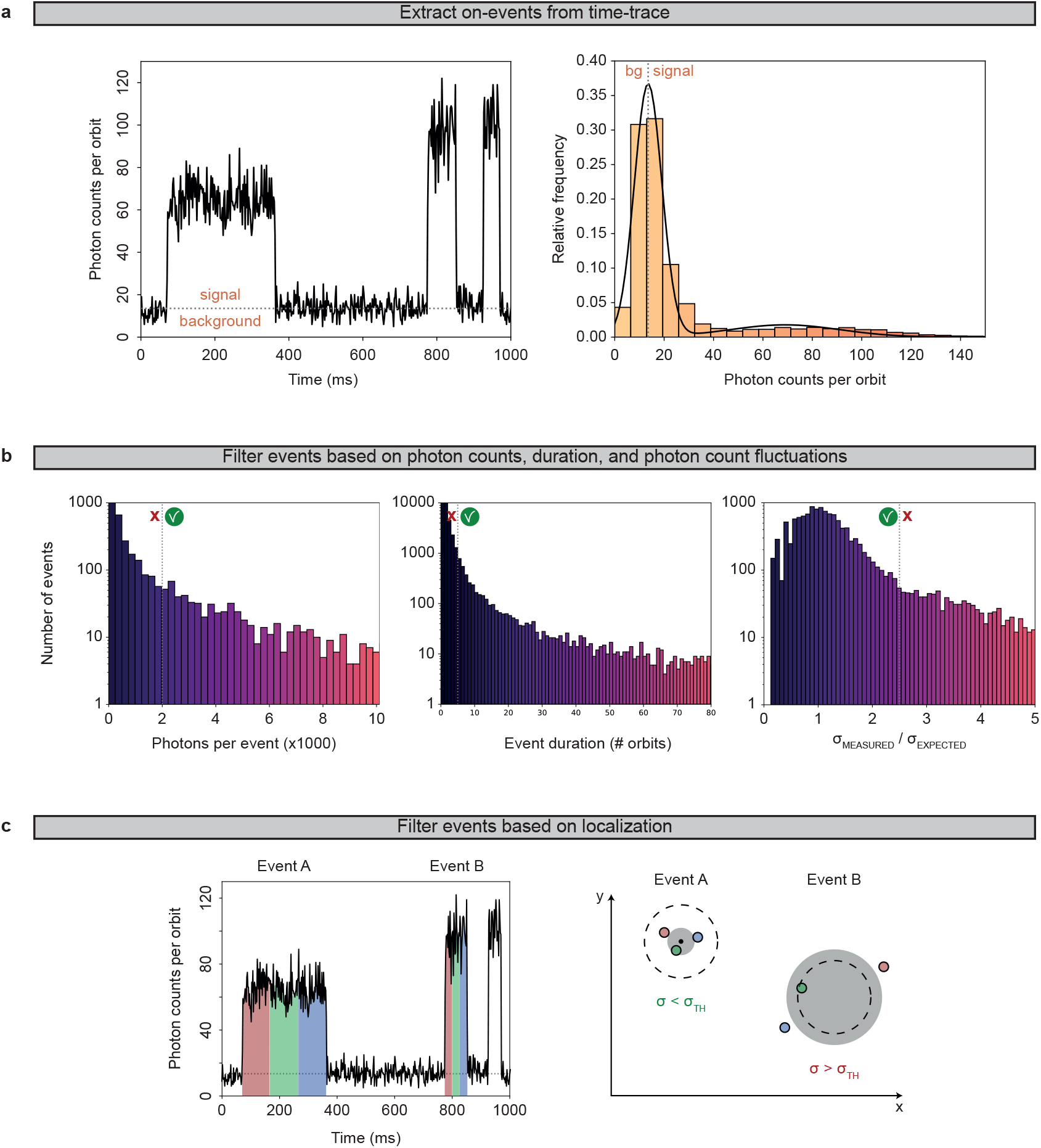
Analysis protocol for ISM-FLUX on DNA-origami measured with DNA-PAINT. (a) Part of the time trace, obtained by summing all photons in all detector channels for all positions on an orbit, and histogram and fit of the full time trace (bg = background). (b) All events pass through three filters, based on the total number of photons (left), the duration (center), and the standard deviation *σ*_measured_ of the count fluctuations within an event (right). (c) Each event that passes all filters is split into three equally long chunks, resulting in three independent localizations, colored in red, green, and blue. If the three localizations are closer to each other than a user-chosen threshold, i.e. *σ < σ*_*T H*_, the event is accepted, otherwise, the event is discarded. The grey circle has a radius of *σ*, the dotted line indicates the threshold. Here, event *A* is accepted, and the three localizations are merged into a single one, obtained by taking the mean coordinates (black dot). Event *B* is discarded.

**Fig. S15.**
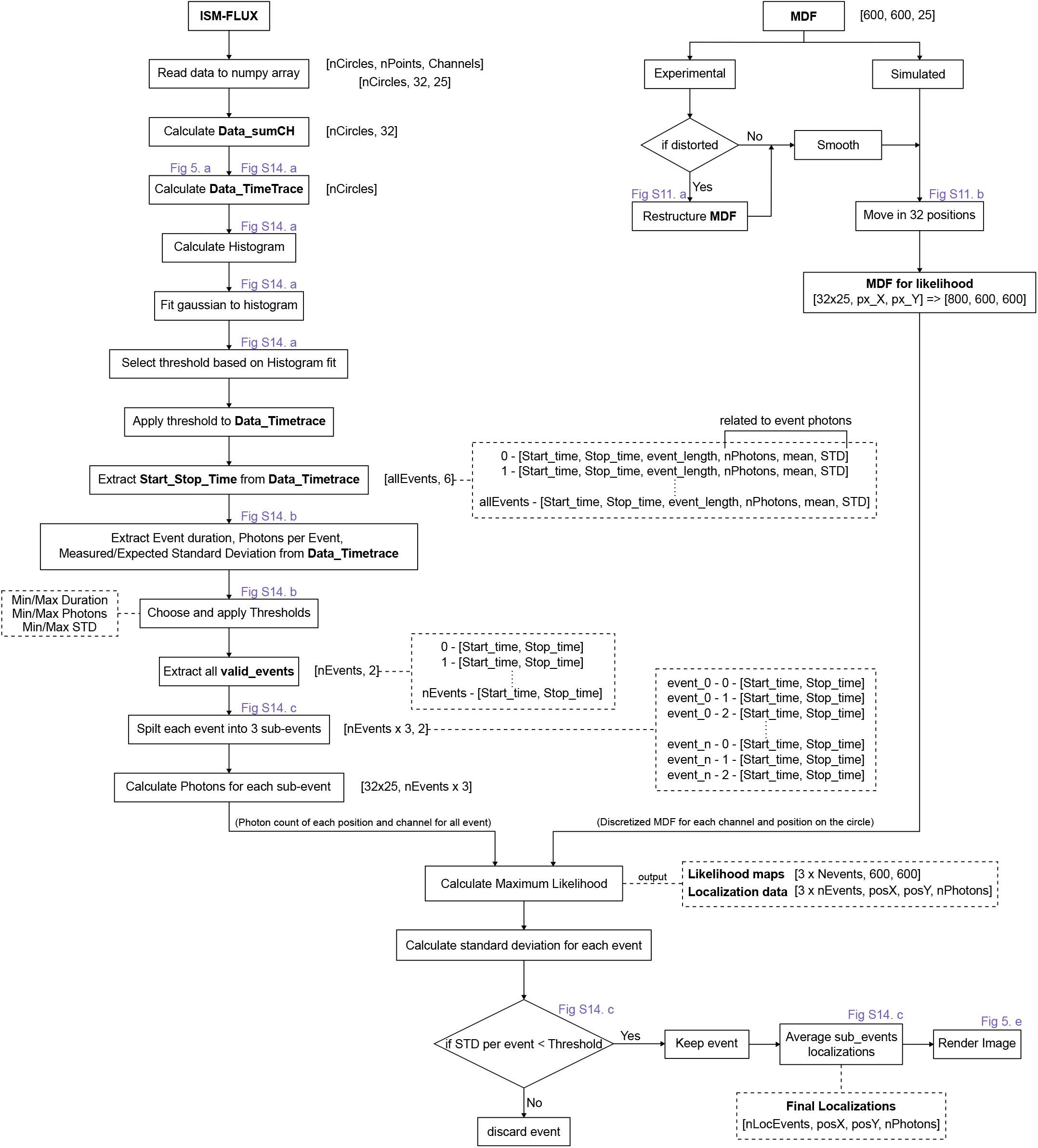
Analysis pipeline for ISM-FLUX data analysis.

